# Protective role of *Pten* downregulation in Huntington’s Disease models

**DOI:** 10.1101/2024.03.28.587147

**Authors:** Nisha, Deepti Thapliyal, Bhavya Gohil, Aninda Sundar Modak, Sanchi Ahuja, N. Tarundas Singh, Bhavani Shankar Sahu, M. Dhruba Singh

## Abstract

Huntington’s disease (HD) is a dominantly inherited neurodegenerative disorder that stems from the expansion of CAG repeats within the coding region of the Huntingtin gene. Currently, there exists no effective therapeutic intervention that can prevent the progression of the disease. Our investigation aims to identify a novel genetic modifier with therapeutic potential. We employ transgenic flies containing *Htt93Q* and *Htt138Q*.mRFP constructs, which encode mutant pathogenic Huntingtin proteins featuring 93 and 138 polyglutamine (Q) repeats, respectively. The resultant mutant protein causes the loss of photoreceptor neurons in the eye and a progressive loss of neuronal tissues in the brain and motor neurons in *Drosophila*. Several findings have demonstrated the association of HD with growth factor signaling defects. Phosphatase and tensin homolog (Pten) have been implicated in the negative regulation of insulin signaling/receptor tyrosine signaling pathway which regulates the growth and survival of cells. In the present study, we downregulated *Pten* and found a significant improvement in morphological phenotypes in the eye, brain, and motor neurons. These findings were further correlated with the enhancement of the functional vision and climbing ability of the flies. We also noted the reduction in both poly(Q) aggregate levels and caspase activity which are involved in the apoptotic pathway. Moreover, we elucidated the protective role of Pten inhibition through the utilization of VO-OHpic (referred to as PTENi). In alignment with the genetic modulation of *Pten*, pharmaceutical inhibition of *Pten* improved the climbing ability of flies and reduced the poly(Q) aggregates and apoptosis levels. A similar reduction in poly(Q) aggregates was observed in the mouse neuronal inducible HD cell line model. Our study illustrates that Pten inhibition is a potential therapeutic approach for HD.

## Introduction

Huntington’s disease (HD) is a dominantly inherited, progressive neurodegenerative disorder characterized by chorea or movement disorder, and cognitive and psychiatric decline [1]. HD is primarily characterized by the expansion of CAG repeats within the first exon of the coding region of the Huntingtin gene (HTT), which results in the formation of toxic polyglutamine [poly(Q)] aggregates that are detrimental to neurons [2]. In healthy individuals, the CAG repeats range from 10 to 35, whereas the detrimental clinical symptoms manifest when the repeat length exceeds 40 (McColgan and Tabrizi, 2018) [3]. The disease typically begins between the ages of 35 to 40 and ultimately leads to mortality within 5 to 20 years following the initial diagnosis [1, 4]. Furthermore, there is an inverse correlation between the CAG repeat length and the age of disease onset. Genetic anticipation where the severity and earlier onset in subsequent generations are evident in HD and juvenile form of HD (JHD) where the disease onset is even before 21 years have been reported [5]. The CAG repeat length in JHD is above 60 however in childhood-onset form the repeat length can exceed more than 80 [5].

The huntingtin gene is predominantly expressed across various tissues and plays a role in neurogenesis [6]. It acts as a protein scaffold, binding to microtubules and actin, and facilitates intracellular trafficking. Additionally, Htt is involved in the transport and regulation of Brain-derived neurotrophic factor (BDNF) [7]. In pathological conditions, the expanded poly(Q) protein has an increased propensity to self-assemble, forming oligomers and subsequently poly(Q) aggregates, leading to a toxic gain of function mutation [3]. In human patients and animal models of poly(Q) diseases, these aggregates disrupt axonal transport, transcription, translation, redox reactions, and protein homeostasis [8-11]. While gain of function results in the disease phenotype, mutations causing loss of function in the Htt protein induce neurodegeneration [12]. Despite significant progress in understanding the genetic and molecular mechanisms underlying HD, there is still much to uncover regarding signaling cascades that could be targeted therapeutically. Presently available therapies primarily focus on symptom management, as neuroprotective treatments aimed at reducing poly(Q) aggregates and halting disease progression remain elusive [4]. This study aims to identify genetic and molecular targets that could have therapeutic applications using the highly tractable *Drosophila* model of HD.

In this study, we investigate the impact of Phosphatase and tensin homolog deleted on chromosome 10 (PTEN), on ameliorating the Huntington’s disease phenotype using the *Drosophila* disease model. Although PTEN was initially identified as a tumor suppressor, its role in the regulation of cellular metabolism has been recently identified [13]. PTEN is a critical component downstream of Receptor Tyrosine Kinase TrkB (including insulin receptor). As a lipid phosphatase, it plays a pivotal role in the dephosphorylation cascade that converts Phosphatidylinositol-triphosphate (PIP3) to Phosphatidylinositol-biphosphate (PIP2). Due to this, PTEN acts as an antagonist to PI3K and thereby enhances PIP2 levels while concurrently reducing phospho-Akt levels, thereby altering the phosphorylation state of numerous downstream proteins involved in cellular proliferation and growth [14-16]. Consequently, this enzymatic function leads to the negative modulation of signaling cascades associated with Insulin and EGFR signaling pathways[15, 17]. In physiological states, these pathways initiate PI3K activity, subsequently resulting in the phosphorylation and activation of AKT, a serine/threonine kinase[18]. Following AKT activation, it phosphorylates multiple downstream proteins such as Gsk3β, mTOR, and TSC, and promotes cellular growth and proliferation[13, 18]. Moreover, it also inactivates pro-apoptotic factors including caspases allowing the survival of cells [19]. Mutation of PTEN increases the level of PIP3 which in turn causes overactivation of AKT which is tumorigenic by induction of abnormal cell growth and proliferation [18]. Furthermore, loss of PTEN function has been shown to increase the protein synthesis by modulation of 4EBP binding protein [20]. *Drosophila* Pten is evolutionary conserved from flies to humans and performs similar functions as its vertebrate counterpart. *Drosophila* Pten also negatively regulates cell growth and cell number, since overexpression of Pten reduces the size of the eye and wing and induces apoptosis[21, 22]. Conversely, the mutation of Pten increases the organ size, including the eye and wing [23, 24]. In the murine models, PTEN downregulation is protective in motor neuron disease and brain injury[25, 26]. In this investigation, we assessed the potential protective effects of Pten downregulation in a *Drosophila* model of Huntington’s disease. We used two transgenic lines of HD models i.e., *Htt93Q* and *Htt138Q*, which express abnormally expanded poly(Q) repeats to solidify our findings[27, 28]. We downregulated Pten in these *Drosophila* models in tissue tissue-specific manner. We observed that Htt93 and *Htt138Q* cause various morphological, functional, and molecular defects. Downregulation of Pten improved the morphological and functional defects Moreover, poly(Q) aggregates and cell death were significantly reduced by knockdown of Pten. These findings from our study propose that inhibition of PTEN is a promising therapeutic target for Huntington’s disease.

## Materials and Methods

### Drosophila Fly stocks

Fly stocks were reared in standard cornmeal media and raised at 25 ± 1°C with a 12h light/dark cycle. *Oregon R+* strain and *UAS-GFP*.*RNAi* (BDSC#9330) was used as a control. The Huntington disease transgenic lines were *UAS-Htt138Q*.*mRFP* [27] and *UAS-Htt93Q* [11, 28]. The following stocks were obtained from Bloomington Drosophila Stock Center (BDSC, USA): *GMR-*Gal4 [29], *UAS-Pten*.*RNAi* (BDSC#8550 & BDSC#25841), *UAS-GFP*.*RNAi* (BDSC#9330), *Or47b-Gal4* (BDSC#9983), *Elav-Gal4* (BDSC#8765).

### External eye and ommatidia imaging

5-20 days adult flies of all genotypes were collected. Flies were fixed on glass slides using nail polish, and one of the eyes faced upward. External eye images were captured using a Stereo zoom bright field microscope (Leica M165 C). Eye size was analyzed using ImageJ software (NIH). To observe the photoreceptors, 2-day-old adult flies of all genotypes were decapitated. The head was mounted and observed using a 60X oil objective under the epifluorescence bright field microscope (Olympus BX53). The aperture was adjusted to allow a narrow light to pass through the rhabdomeres. The number of photoreceptors in each genotype was quantified in ImageJ software (NIH).

### Immunostaining

For immunostaining, eye discs and adult brains were dissected in Phosphate buffer saline (PBS), fixed for 20 minutes with 4% paraformaldehyde, and washed three times with PBST (with 0.1% Triton X-100). Tissues were blocked with PBST with 0.1% BSA solution and incubated with primary antibody overnight at 4°C. Primary antibodies used are, anti-Dcp-1 (1:100; 9578; Cell Signaling Technology, USA), anti-Dlg1 (1:100, 4F3, DSHB, USA), anti-GFP (1:100, 12A6, DSHB, USA), anti-FasII (1:25, 1D4, DSHB, USA), mouse anti-Futsch (1:100, 22C10, DSHB, USA). Tissues were then incubated in secondary antibody for 2 hours with gentle shaking in the dark. The secondary antibodies used were Alexa 488 goat anti-rabbit (A11034), Alexa 488 goat anti-mouse (A28175), Cy3 goat anti-mouse (A10521), and Alexa 647 goat anti-mouse (A31571). The tissues were washed in PBST and counterstained with DAPI (5μg/ml, D3571, Invitrogen, USA) then mounted using anti-fade media (P36934, Molecular probes, USA).

In the quantification of Dcp-1 staining at the larval eye disc stage, the area below the morphogenetic furrow was selected in each sample and Dcp-1 positive puncta were counted by using ImageJ software after applying similar threshold values in samples. The mean values with ±SD were plotted in graphical form. In the case of pupal eye tissue, an ROI was drawn randomly with a fixed area in each sample tissue (i.e., 100μm^2^). Similar parameters were followed for the entire sample.

### Histology

For histological sections, 2-day-old fly heads were decapitated in Phosphate-buffered saline (PBS) and fixed for 90 minutes in 4% paraformaldehyde. Subsequently, adult heads were then dehydrated through a series of increasing ethanol concentrations in PBS for about 40 minutes each in 50%, 70%, 95%, and 100% ethanol solutions. It was followed by xylene washes with overnight incubation at RT in 1:1 xylene and Paraffin wax. The next day wax washes were given 4 times for 2 hours at 60°C. Later the fly heads were embedded in paraffin wax in proper orientation. Using a microtome (YSI 055, Manual Rotary microtome, Yorco, India), serial frontal 16µm sections of adult heads were prepared on gelatin-coated slides. Slides were then dewaxed in xylene and rehydrated in decreasing ethanol concentrations in PBS for about 5 minutes each in 100%, 95%, 70%, and 50% ethanol. Finally, tissues were stained in 0.01% Toluidine blue solution (Cat no. SRL, 22134) and mounted in DPX medium. The images were captured at 10X.

### Adult NMJ preparation

To prepare samples of adult thorax Neuromuscular Junction (NMJ), 2 or 5-day-old adult flies were selected. The Dorsal Longitudinal Muscles (DLMs) were dissected after removing the head and abdomen from the thorax. Subsequently, the samples were fixed in a solution of 4% paraformaldehyde in PBS for 20 minutes and then washed four times with PBT at room temperature (RT). The thoraces were flash-frozen with liquid nitrogen and bisected down the midline in ice-cold PBS. The tissues were then incubated in a blocking buffer containing PBST with 0.1% BSA for a minimum of 2 hours at Room temperature, as described earlier [30]. For immunostaining, the following primary antibodies were used for overnight incubation at 4°C.e., anti-Futsch (1:100, DHSB, USA). Following primary antibody incubation, the tissues were washed four times in PBST for 10 minutes each and then incubated in secondary antibodies. The appropriate secondary antibodies were employed for 2 hours incubation at room temperature. The list of secondary antibodies used are, Alexa-488 (Molecular Probes) at a dilution of 1:200, Cy3-conjugated anti-HRP (cat no. 123-165-021, Jackson immune research) at a dilution of 1:500, and FITC-conjugated anti-HRP (cat no. AB 2338965, Jackson immune research) at a dilution of 1:200. Following the secondary antibody incubation, samples underwent washing with PBST four times and then subsequently mounted on a glass slide using Vectashield mounting media (Vector Laboratories).

Images for DLM synaptic morphology were acquired using a 63X objective oil lens (Numerical aperture 1.4). Z-stacks were generated at a consistent tissue depth of 30 slices, with a 0.5μm interval. The images were taken starting from the point where Horse Radish Peroxidase (HRP) staining initially appeared at muscle fiber III, indicated by the white box in Figure 3B. In total, 10 images were captured per experimental condition using identical parameters. The images were processed as Max Intensity Projections (MIP) using Fiji software [31]. To measure synaptic morphology, total neurite length (in µM) and branch numbers were determined by tracing HRP staining, facilitated by the updated Simple Neurite Tracer (SNT) Plug-in [32, 33]. The analysis was conducted using the Skeletonize 3D Plug-in of Fiji [31, 34].

### Quantitative RT-PCR

RNA extraction was done from age-matched adult *Drosophila* heads (n=60) using Trizol (Cat no. T9424, Sigma-Aldrich, USA) reagent as per manufacturer protocol. The cDNA was synthesized from 1μg of total RNA using the kit (cat no. 1708891, Bio-Rad) as described earlier [35]. Real-time PCR was performed using qPCR SYBR Green master mix (Bio-Rad) in the CFX-96 Bio-Rad PCR machine (Bio-Rad, USA). The corresponding primer sequences used are:

*rp49 (F)* : ATGCTAAGCTGTCGCACAA

*rp49(R)* : TTGTGCACCAGGAACT TCTT

*Pten(F)* : CAGTTTCCGGCGATGTAAAA

*Pten(R)* : ACATCATCGATTTCTGATTTGC

### Lifespan assay

Around 100 freshly eclosed adult flies were collected and 10 flies were transferred per vial which contained fly media. Flies were transferred to fresh vials every alternate day after assessing the survival in each vial until all flies had died. The mean lifespan of flies was calculated, and the Kaplan-Meier plot was generated using the online OASIS tool and GraphPad Prism.

### Negative geotaxis assay

Negative geotaxis was conducted to assess the locomotor and motor ability of the flies [36]. About 100 flies per genotype were collected in batches of 10. Subsequently, ten flies were introduced to a vertical column and were given five minutes to acclimatize to the new environment. The vials were then gently tapped to encourage all the flies to descend to the bottom of the vial. The number of flies crossing the 8 cm or 4 cm mark in the vertical column in 10 seconds was recorded. The assessment was conducted at the age of 5-, 10-, 15- and 20-days post eclosion. The results were analyzed to measure the climbing ability of each genotype with age.

### Phototaxis assay

The Phototaxis assay was performed to evaluate the alteration of functional vision. About 100 flies from each genotype were collected. Ten flies were introduced into a Y-maze made of glass; one arm of the glass was covered with black film to prevent light from entering, and the other arm light was illuminated by an LED bulb. The flies were collected after gentle tapping from either arm of the Y-maze after 10 seconds and the number was counted in dark and light condition. The experiments were performed at the age of day 5 and day 20 post-eclosion.

### Drug feeding

For the inhibition of Pten, VO-OHpic trihydrate was used (Santa Cruz, Sc-216061A). The stock solution was prepared by dissolution in DMSO [37]. To identify the optimum working concentration of the drug, cornmeal agar food media were prepared with different concentrations of the drug ranging from 10μM to 100μM. The final concentration of the drug 50µM/ml was used in further experiments. Protocol reported by Deshpande et al., 2014 [38], was adapted for drug feeding. In brief, eggs from the appropriate genotypes were collected overnight on agar plates, and corresponding first instar larvae were transferred into normal food containing DMSO and drug-food containing VO-OHpic (50μM). Further experiments were performed at the larval and adult stages. Results were plotted to see the difference in Htt aggregates and behavior in drug-fed and DMSO-fed flies.

### Cell culture and treatments

Mouse neuroblastoma N2a cells with Ponesterone A (Enzo, ALX-370-014-M001) inducible truncated N-terminal Huntingtin gene with 150 CAG repeats attached to EGFP (HD 150Q) were a kind gift from Dr Nihar Ranjan Jana Lab, NBRC. The cells were grown in DMEM Glutamax, supplemented with 10% FBS, 1% antibiotic-antimycotic along with 0.4mg/ml of G418 and 0.4mg/ml of Zeocin (Invitrogen) at 37° C and 5% CO2 in a humidified incubator. Early Passage cells (1-10) were used for all the experiments. Mouse N2a cells were induced with Ponesterone A, adding 20nM and 50nM of VO-OHpic (PTENi) to each condition. After 1 day of induction and treatment with VO-OHic, the cells were harvested and lysed for various experiments such as Immunoblot and Fluorescence Imaging.

### Protein extraction and Western Blot

Following the completion of the initial treatment, cells were lysed with RIPA lysis buffer (50 Tris, pH 7.4, 1mM EDTA, 150 NaCl, 1% Triton -X, 0.1% SDS, 1% Sodium Deoxycholate with 1mM phenylmethylsulphonyl fluoride, 1mM NaF, 2mM Na2VO5, 20mM Na4P2O7 as a phosphatase inhibitor cocktail (Complete, 11873580001) at 4°C. Afterwards, the cell lysates were collected in a small centrifuge tube and centrifuged at 14,000 g for 15 minutes and the supernatant was collected in a fresh centrifuge tube. The protein content of the samples was estimated, following Takara’s BCA kit manufacturer’s protocol. Following protein estimation, 20μg protein was loaded on SDS-PAGE to separate proteins based on their molecular weight and transferred onto nitrocellulose membrane (MDI, SCNG8101XXXX101-4). The Nitrocellulose membrane was blocked with 5% skimmed milk or 3% BSA (only for phosphoprotein) for 2 hours at room temperature. After blocking, the membranes were incubated with primary antibodies against GFP (1:10000) (Proteintech, 50430-2) and GAPDH (1:40000) (Proteintech, 60004-1-IG) overnight at 4°C. After 12-14 hours of incubation, the membranes were washed with Tris-buffered saline with 0.1% Tween 20 (TBST) and incubated with HRP-conjugated goat antimouse (1:10000) and goat anti-rabbit (1:10000) secondary antibodies. Images were captured using the UVitec Mini HD9 gel imaging system. Densitometry quantification was done using Image Lab (6.0.1)

### Fluorescence imaging

2*10^4^ cells were seeded on each 15 mm coverslip, and the cells were treated with the VO-OHpic (PTENi) (50nM) by the above-discussed protocol. After treatment, cells were gently washed with 1x PBS twice and then fixed using 4% Paraformaldehyde (PFA) for 20 minutes at room temperature. Following fixation, cells were washed gently with 1x PBS twice. Coverslips were mounted using DAPI containing mounting media, and images were captured using 20x of Zeiss-Apotome Microscope.

## Statistical analysis

All graphs and statistical analysis were performed using Prism GraphPad software (GraphPad 6.0). Data represents mean, standard deviation (SD), and n values. As per the experimental requirement based on single or multiple parameter analysis, data were subjected to a two-tailed unpaired t-test, one-way analysis of variance (ANOVA), or two-way ANOVA. A p-value less than 0.05 was considered significant. For survival assay, the survivorship curve uses the Kaplan-Meier approach.

## Results

### Downregulation of Pten improves the eye phenotype of the *Drosophila* model of HD

We used the *Drosophila* models of HD expressing mutant Htt protein with *93Q* and *138Q* repeats [27, 28]. The 138Q is tagged with RFP, which allows the observation of poly(Q) aggregates [27]. *GMR-Gal4* was used to express the mutant form of Htt transgene in the eyes of developing and adult flies. Targeted expression of human *Htt93Q* using eye-specific *GMR-Gal4* driver exhibited a strong neurodegenerative phenotype characterized by loss of pigmentation and reduced eye size at 20 days of age (Fig. 1C) in comparison to control flies (Fig. 1A, B). 5-day old flies did not show any such eye phenotypic defects (Fig. S1C) but developed the phenotype upon aging (Fig. 1C). However, *Htt138Q* repeats in the fly eye caused a rough eye phenotype even at 5 days (Fig. S1E) which gets more intensified upon aging (20-day) (see Fig. 1E and S1E). Knocking down of *Pten* in this context improved the roughness of the external eye morphology and reinstated the amount of pigmentation of photoreceptor cells (Fig. 1D and F). Downregulation of *Pten* exhibited no change in the adult eye phenotype at both 5 days and 20 days (Fig. 1B and S1B). Quantification data showed a significant reduction in eye size in both 93Q and 138Q expressing flies (Fig. 1S). Knockdown of *Pten* significantly improved the eye size (Fig. 1S). To further illustrate the improvement in ommatidial structure, 40-hour-old pupal eyes were stained with a Disc large (Dlg). Compared to the wild type no ommatidial defects were observed in *Htt93Q* pupal tissue (Fig. S1G-I). However, flies expressing *Htt138Q*.*mRFP* exhibited gross morphological defects in primary, secondary, and tertiary cells, cone cells, and abnormal bristle lattice (arrow in Fig. S1K). These defects were partially suppressed following the downregulation of *Pten* in *Htt138Q*.*mRFP* expressing pupal eye tissues (Fig. S1L). These findings strongly indicate that the downregulation of *Pten* effectively curbs the manifestation of phenotypic abnormalities from the initial stages of the disease pathogenesis. We used 2 different transgenic RNAi of *Pten* (BDSC #8550 and BDSC #25841) and found consistent results (Fig. S1M, N). Real-time PCR analysis revealed that both RNAi-mediated downregulation of *Pten* lines (BDSC #8550 and BDSC #25841) in the brain tissue reduced the expression by 0.5 and 0.6 fold respectively.

**Fig. 1.**
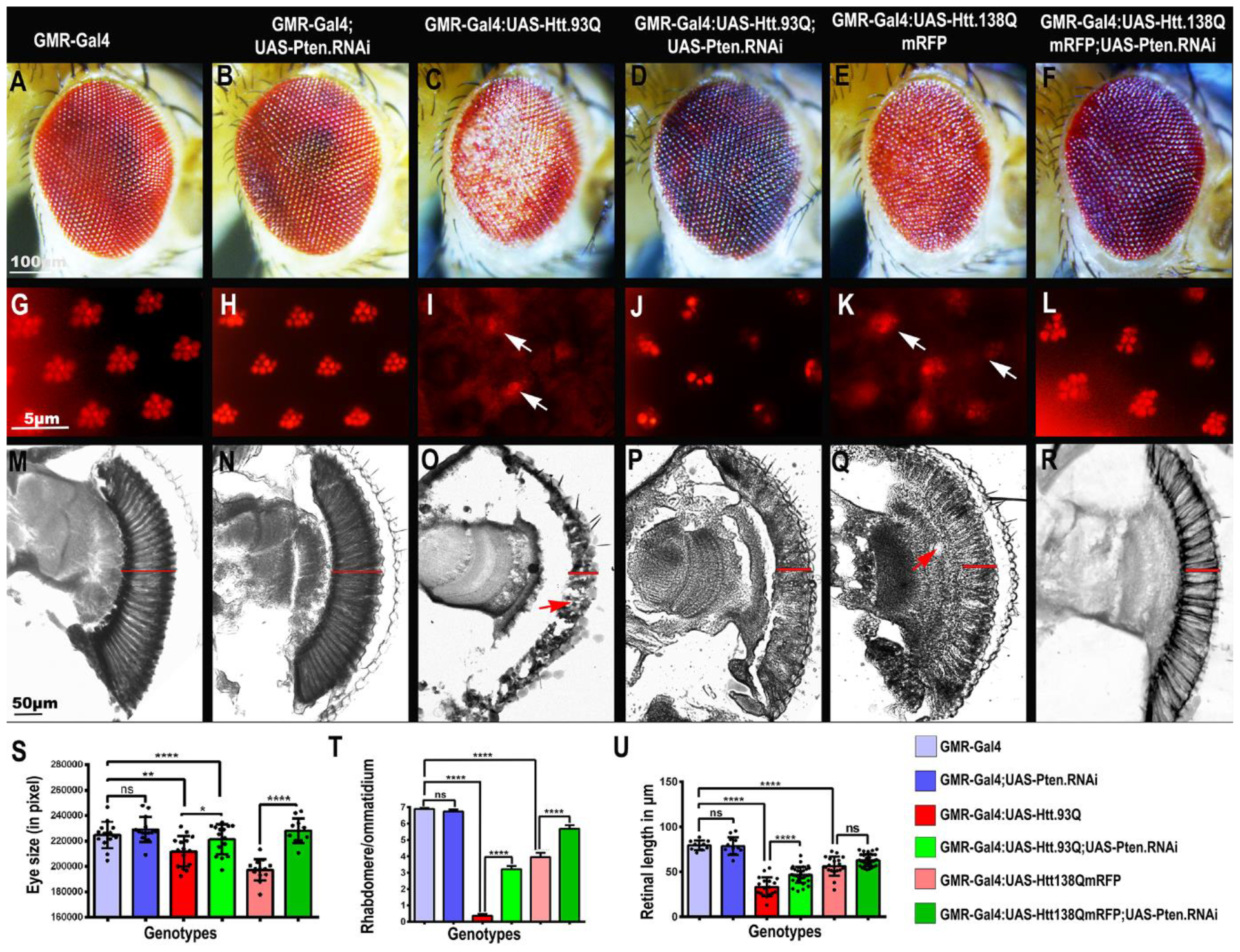
Knockdown of Pten improves the *Htt93Q* and *Htt138Q-induced* eye degeneration. (A-F) Bright-field images of the 20-day-old adult eye. (A) *GMR-Gal4* (B) *GMR-Gal4;UAS-Pten*.*RNAi* (C, E) Overexpression of *Htt93Q* and *Htt138Q*.*mRFP* resulted in decreased eye size and depigmentation. (D, F) Downregulation of *Pten* in *Htt93Q* and *Htt138Q*.*mRFP* background improved the eye pigmentation and reduced the rough eye phenotype in both the diseased flies, respectively. (G-L) Deep pseudo pupil images of 2-day-old adult eye (G-H) *GMR-Gal4* and *UAS-Pten*.*RNAi* have 7 photoreceptor cells whereas (I, K) *Htt93Q* and *Htt138Q*.*mRFP* flies showed degeneration of photoreceptor cells (arrows). (J, L) Downregulation of *Pten* improved the number of photoreceptors. (M-R) Horizontal sections of the eye retina of 2-day-old flies. (M-N) *GMR* and *Pten*.*RNAi* control flies (O) Retinal tissue is degraded due to the expression of *Htt93Q*. (P) Downregulation of *Pten* restored retinal projections. (Q) *Htt138Q* flies showed degeneration and vacuoles in the retinal region. (R) *UAS-Htt138Q*.*mRFP; UAS-Pten*.*RNAi* showed improved retinal projections (S) Bar graph showing the difference in eye size (T) Bar graph showing the number of ommatidium (U) Bar graph indicating the retinal thickness. Student’s t-test; Mean ± SE (ns= non-significant, ns P>0.05, *P<0.05, **P<0.01, ****P<0.0001). Scale bars: (A-F) 100µm, (G-L) 5 µm, (M-R) 50µm.

However, diseased flies (*Elav-Gal4/UAS-Htt93Q*) showed a non-significant upsurge in *Pten* transcript level (see Fig. S1O, n=3). In fact, in rescue flies it appears due to this upsurge *Pten* transcript was reduced only up to 0.3-fold, which suggests mutant Htt may play a significant role in regulating *Pten* transcript level.

Additionally, the pseudo-pupil technique was utilized to observe the enhancement in photoreceptor neurons in adult flies. The wild type and *Pten*.*RNAi-expressing* flies displayed 7 visible photoreceptors; however, *Htt93Q* and *Htt138Q* expressing diseased flies revealed substantial degeneration of photoreceptor cells (compare Fig. 1G to I and K). An average of only 1 to 4 photoreceptor(s) was evident in each ommatidium (arrows in Fig. 1I and K) of flies expressing *Htt93Q* and *Htt138Q*. Interestingly, RNAi-mediated downregulation of *Pten* in *Htt93Q* and *Htt138Q* expressing tissues constrained the degeneration of the photoreceptor cells, and each ommatidium displayed an average of 3 to 6 photoreceptors (Fig. 1J and L).

To further illustrate the improvement in retinal tissue, the adult eye tissue was fixed, and tangential sections were generated following paraffin molding. The tangential section of the eye shows retinal projections from the ommatidia into the lamella of the optic lobe in control (*GMR-Gal4*) and *Pten-RNAi* flies (Fig. 1M and N). Flies expressing *Htt93Q* and *Htt138Q* exhibited loss of retinal tissue (Fig.1O and Q). In *Htt93Q*, only a thin layer of retinal tissue was observed below the ommatidial cells. Knockdown of *Pten* mitigated the retinal tissue degeneration, and improved retinal length (compare Fig. 1O and P). Likewise, retinal tissue showed degeneration in *Htt138Q*, which was also ameliorated by the knockdown of *Pten* (Fig. 1Q and R). Furthermore, we checked for the improvement of functional vision by introducing the flies into the Y-maze and counting the number of flies that went to the dark or illuminated arm of the Y-maze. For both male and female flies of *Htt93Q*, the phototaxis behavior towards the illuminated arm was reduced. Interestingly, the knockdown of Pten improved the percentage of flies moving to the lighted arm (Fig. S1P). The above results showed that reduced *Pten* expressio*n* is protective against mutant Htt-induced tissue degeneration.

### Knockdown of Pten suppresses the degeneration of neuronal tissues in the brain and consequently exhibits enhanced survival

Since Htt is predominantly expressed in the brain, we sought to determine whether the knockdown of *Pten* could elicit a similar protective function in the brain and motor neurons as seen in the eye model. To achieve this, we employed the pan-neuronal driver, *Elav-Gal4*, to induce the expression of mutant Htt protein in neuronal tissues. We focused our attention on the *Drosophila* mushroom body, a neuronal structure involved in olfactory memory and learning, consisting of Kenyon cells and their axonal projections which form the α, β, and γ lobes [39, 40]. Staining with Fas-II revealed the presence of α, β, and γ lobes in the control adult fly brain (Fig. 2A-B). However, flies overexpressing *Htt93Q* showed progressive degeneration of mushroom body structures in 5 and 20-day-old flies (Fig. 2C and S2C). Visually, we categorized the Mushroom body structure into Mild, Moderate, and Severe forms. Notably, moderate, and severely damaged mushroom bodies were higher in *Htt93Q* flies and Mild and moderately damaged mushroom bodies were higher in flies co-expressing *Pten*.*RNAi* (Fig. 2D, E). Furthermore, our investigation revealed that *Htt138Q* caused more severe mushroom body degeneration even at 10 days and *Pten* knockdown mitigated the neuronal loss (Fig. S2E-L, M).

**Fig. 2.**
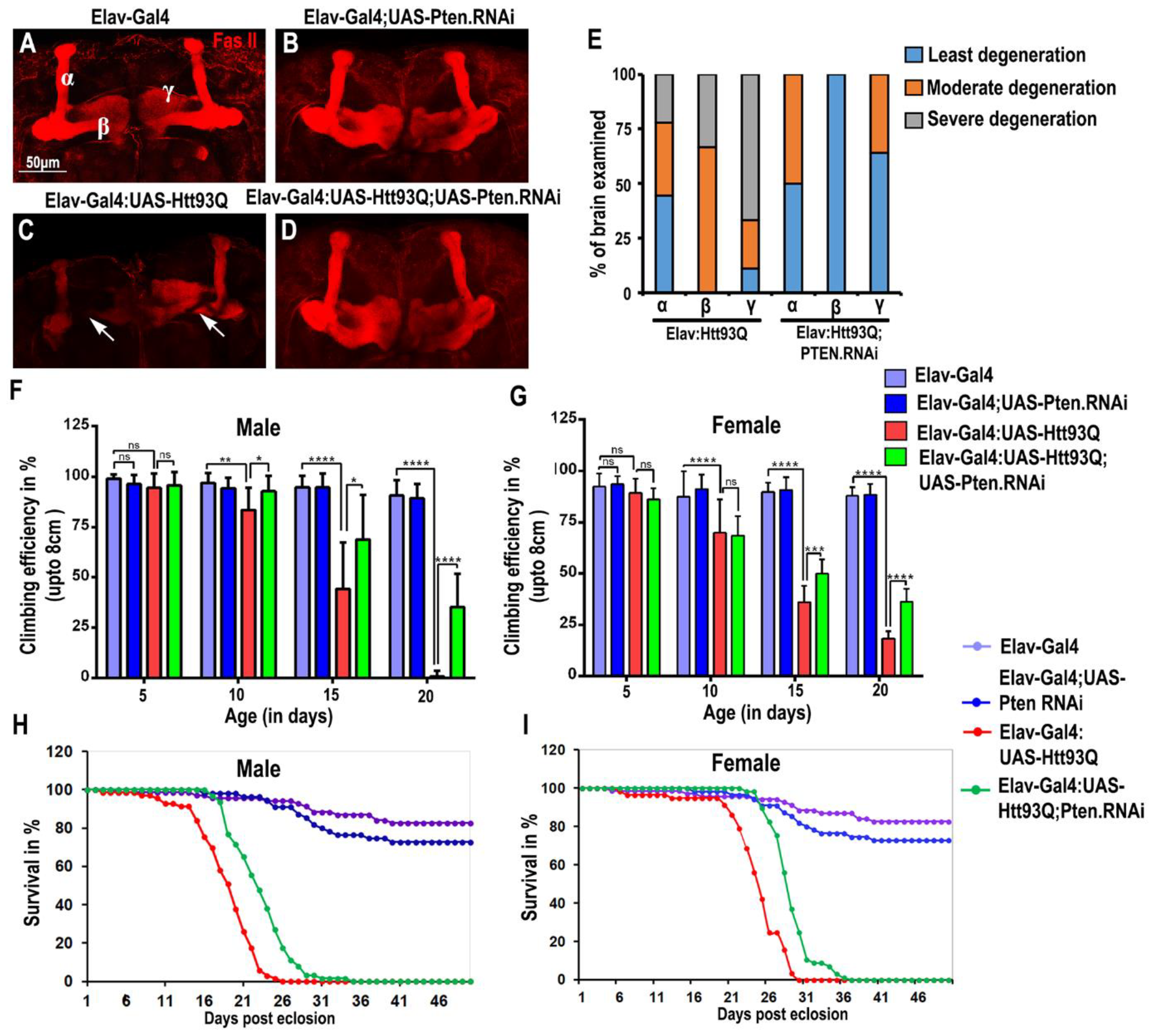
Pten knockdown suppresses the *Htt93Q*-mediated brain degeneration and improves climbing behavior and Longevity. (A-D) 20 days old adult brain stained with Fas-II (red) antibody. (A, B) α, β, and γ lobes of the mushroom body are intact in control flies. (C) *Htt93Q* caused mushroom body degeneration which exhibited the absence of γ lobe and degeneration of α, β lobe. (D) Downregulation of *Pten* restored the three α, β, and γ lobe structures. (E) Graphical representation comparing the percentage of degeneration of α, β, and γ lobes of the mushroom body. (F & G) The bar graph represents the climbing ability of 5, 10, 15, and 20-day-old flies. *Pten*.*RNAi* significantly ameliorated the locomotor ability of *Htt93Q* from 15 days onwards. (H-I) The survival curve showed improvement in the longevity of the flies due to the knockdown of *Pten*. Survival curves were generated by the Kaplan-Meier method and statistical significance was determined by the log-rank test (P<0.05). Error bars represent mean± SE, one-way ANOVA test (α=0.05) (ns= non-significant, *P<0.05, **P<0.01, ***P<0.001, ****P<0.0001). Red: FasII. Scale bar (A-D): 50µm.

To ascertain whether the improvement in brain degeneration correlates with the functional abilities of the flies, we evaluated their climbing ability. Flies that successfully climbed the 8 cm mark in a plastic vial were quantified to assess their motor skills as shown earlier [36]. Interestingly, the climbing ability in both male and female *Htt93Q* flies remained only mildly affected for up to 10 days (Fig. 2F and G,). However, at 15 days, a significant decline in climbing ability was observed compared to control flies, and by 20 days, the motor skills of the flies were severely impaired, with only a few managing to reach the 8 cm mark (Fig. 2F and G). In contrast, significant improvement in climbing was noted in both the males and females when *Pten* was downregulated in *Htt93Q*-expressing flies (Fig. 2F and G). This demonstrates that reduced expression of *Pten* restricts the *Htt93Q* and *Htt138Q-mediated* cellular degeneration and confers functional rescue. Additionally, we assessed the fitness of the flies by monitoring their longevity. We collected around 100 flies for each genotype and mortality was recorded daily to determine their lifespan. Subsequently, the knockdown of *Pten* led to a marked improvement in longevity for both male and female flies in the *Htt93Q* disease background (Fig. 2H, I). Similar observations were made in flies expressing *Htt138Q* (Fig. S2N). These findings underscore the potential of *Pten* knockdown to alleviate neurodegeneration and improve these flies’ overall health and longevity in the context of mutant Httinduced degenerative conditions.

### Motor neuron degeneration improved by knockdown of *Pten*

Expression of mutant Huntington protein with abnormal poly(Q) repeats caused motor neuron defects [41]. Henceforth, the structural integrity of Neuro-Muscular Junctions (NMJs) was investigated in Elav-Gal4/UAS-*Htt93Q* and Elav-Gal4/UAS-*Htt138Q*.mRFP diseased flies. We used the Horse Radish Peroxidase (HRP) antibody to mark the motor neurons in a 2–5 days adult thorax to observe any structural defects. The schematic diagram shows all Dorso-lateral muscles (DLMs) and motor neurons innervating the muscles (Fig. 3A and B). We captured the confocal images of the NMJ from similar regions of DLMs. The control Elav-Gal4 and UAS-Pten.RNAi showed well-arranged NMJs with secondary and tertiary branches (Fig. 3C, D and G, H). In contrast, *Htt93Q* and *Htt138Q* expressing transgenic flies showed reduced branching patterns and severe degeneration of secondary and tertiary branches of motor neurons (Fig. 3E, and I). Thus, the structural deterioration of NMJs in diseased flies was consistent with the behavioral climbing impairment. Interestingly, the downregulation of Pten showed a significant restoration of motor neuron morphology in branching patterns and their arrangements (compare arrowhead in Fig. 3E-F, I-J). Quantification analysis showed a significant reduction in branching and triple points in diseased flies compared to the control, which was improved in Elav-Gal4, UAS-*Htt93Q*/UAS-Pten.RNAi expressing flies (Fig. 3K-L, n=10). Similar analyses were also done for *Htt138Q* flies which also confirmed a gross level of degeneration of the branching pattern of motor neurons in selected DLMs, which was significantly reinstated with down-regulation of Pten in disease background (Fig. 3I-J, M-N). Similarly, anti-Futsch (22C10) staining showed branching defects of motor neurons in *Htt138Q* and improvement by coexpression of Pten.RNAi (compare Fig. S3C and D, also see E and F). These observations suggest that the knockdown of Pten has a protective role and restricts motor neuron degeneration in mutant Htt-expressing flies.

**Fig. 3:**
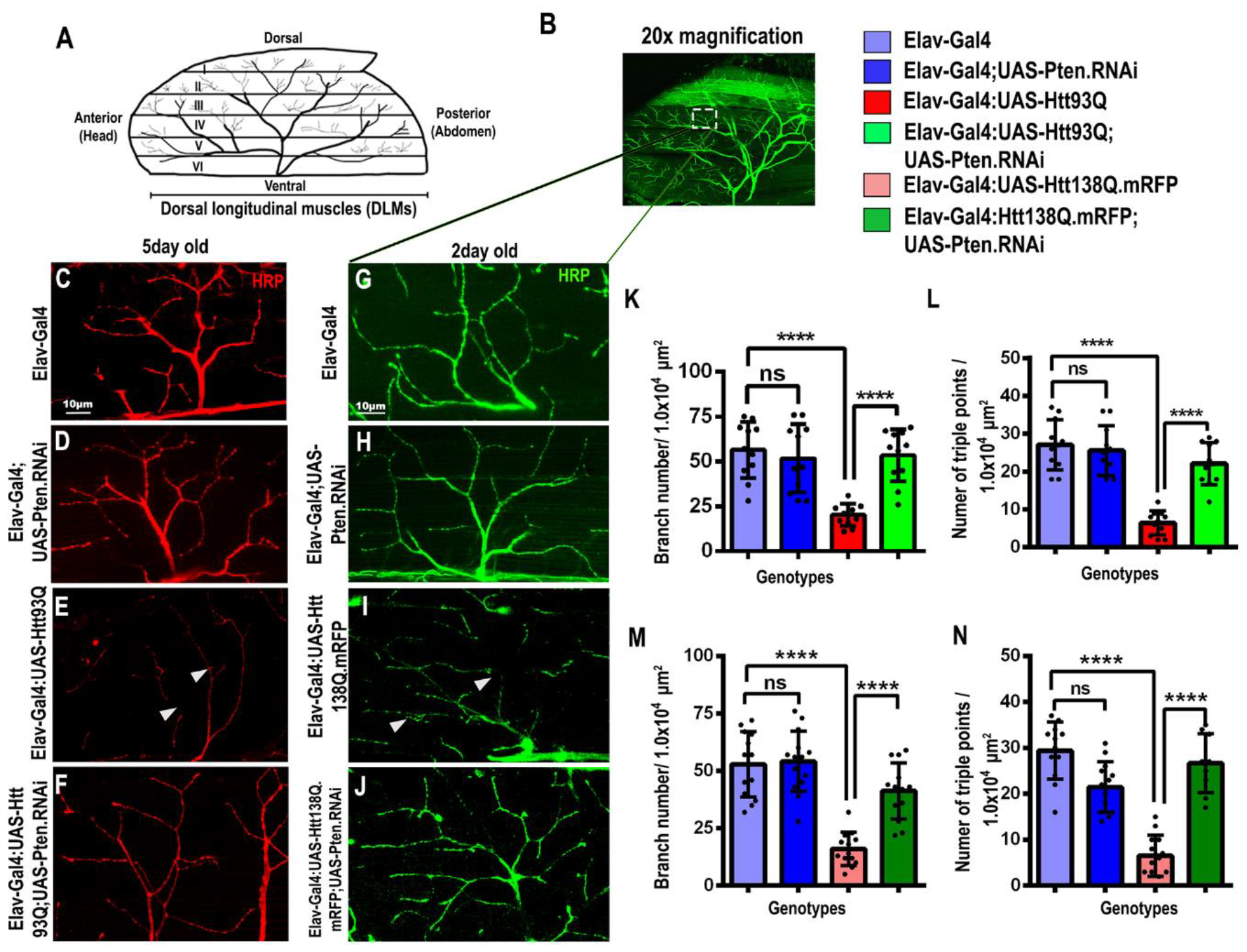
Pten knockdown rescued the mutant Htt-mediated degeneration of synaptic structure in thoracic muscles. (A) Schematic representation of dorsal longitudinal muscle (DLM) along with the typical representation of neuromuscular junctions (NMJs) of hemithorax of the adult fly. (B) 20X magnified image of muscle segments and NMJs which were stained with horseradish peroxidase (HRP). (C-J) HRP stained NMJs of 5-day-old flies. (C and G) *Elav-Gal4*; (D and H) *UAS-Pten*.*RNAi* shows well-arranged branching of motor neurons in DLMs (E and I) Expression of *Htt93Q* and *Htt138Q* resulted in degeneration and loss of branching of motor neurons (Arrows represent the point of synaptic degeneration. (Fand J) Pten.RNAi restored the branching pattern in diseased *Htt93Q* and *Htt138Q* flies. (K-L) The bar graph shows the quantification of the branching number and triple points in the fixed ROI area in each *Htt93Q* fly. (M-N) Similar results were seen in *Htt138Q* flies. Student’s t-test, error bars represent mean± SE (ns= non-significant, ns P>0.05, **P<0.01, ***P<0.001, ****P<0.0001). Green: HRP (C-F), Red: HRP (G-J). Scale bar (C-J): 10µm.

### Pten knockdown reduces the poly(Q) aggregates

The accumulation of protein aggregates in the HD brain is a prominent hallmark of HD pathogenesis and to date, most suppressors of poly(Q) toxicity have been shown to aid in clearing these protein aggregates. We next investigated whether the reduction in poly(Q) phenotypes in flies is due to changes in the level of poly(Q) aggregates. First, we checked the status of aggregates in adult eyes. The *Htt138Q* is tagged with RFP which allows observation of the poly(Q) aggregates under the fluorescence microscope (Fig. 4A-D).

**Fig. 4.**
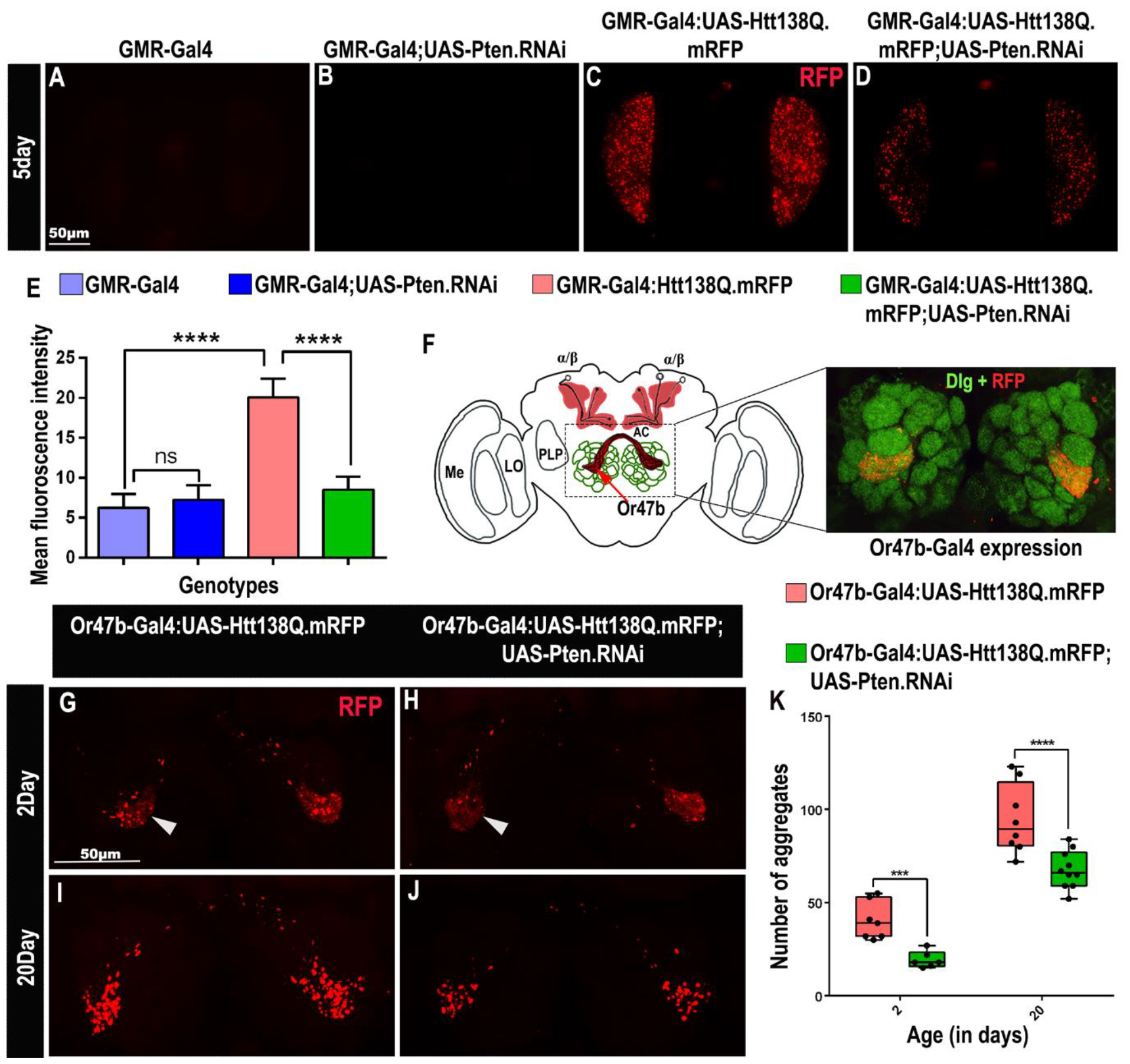
Pten knockdown reduces Htt aggregates in the eye and antennal lobe. (A-D) Adult eye images. (C) *HttQ138*.*mRFP* flies showed aggregates in 5-day-old adult flies. (D) Downregulation of *Pten* significantly reduces the level of Huntingtin protein aggregates. (E) Bar graph showing mean fluorescence intensity of Htt protein aggregates. (F) Schematic representation of the *Drosophila* brain showing the antennal lobe and its expression pattern of odorant receptor 47b specific *Or47b-GAL4*. Antenatal lobe stained by Disc large (Dlg-1) and *Htt138Q*.*mRFP* aggregates. (G-H) 2-day old *Htt138Q*.*mRFP-expressing* flies showed poly(Q) aggregates which were significantly reduced in number by the knockdown of Pten. (I-J) The level of poly(Q) aggregates gets more aggravated in 20-day-old flies which was significantly reduced upon downregulation of *Pten*. (K) Box plot showing poly (Q) aggregates at 2-day and 20-day-old flies of each genotype. Error bars represent mean± SE (ns= non-significant, ns P>0.05, ***P<0.001, ****P<0.0001). Green: Dlg1; Red: *Htt138Q*.RFP. Scale bar: 50µm.

5 days adult flies of desired genotypes were decapitated, and the entire head was mounted on a glass slide and observed under fluorescence microscopes. In comparison to *GMR-Gal4* and *GMR-Gal4; Pten*.*RNAi* (Fig. 4A B) which showed no RFP level, while *Htt138Q*.*mRFP-expressing* flies exhibited a significantly enhanced level of fluorescence intensity in the form of puncta which was significantly reduced by the expression of *Pten*.*RNAi* (compare Fig. 5C and D). To restrict the expression of mutant Htt protein adult fly and observe the changes in the formation of aggregates progressively, we employed *Or47b-Gal4* which is expressed in the specific antennal lobe. The expression of *Or47b-Gal4* was minimal during the early stage, it started expressing after the flies were eclosed from the pupal stage [42]. At the age of 2 days post eclosion, as compared to the controls, the *Htt138Q* expressing flies showed the formation of aggregates (Fig. 4G, S4A-B). However, in *Pten*.*RNAi* expressing flies the diffuse Htt expression was observed, and the number of aggregates was significantly reduced (Fig. 4H). At 20 days, as compared to the control the number of aggregates substantially increased in *Htt138Q* expressing flies while knockdown of *Pten* significantly restricted the enhancement of aggregate formation even in 20-day-old flies (Fig. 4I -K, S4C-D).

**Fig. 5.**
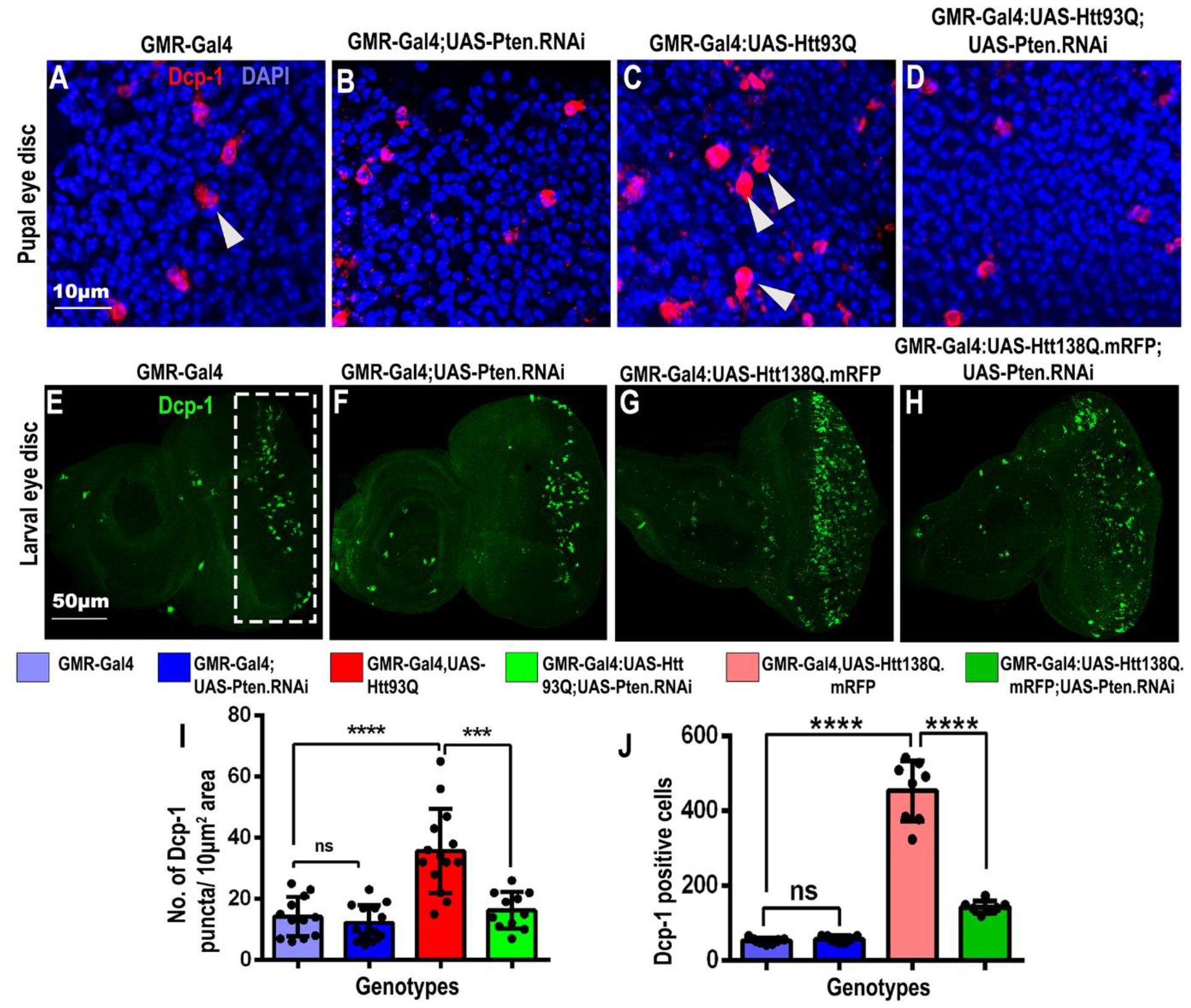
Pten knockdown suppresses cell death induced by mutant Htt protein. (A-J) Larval and pupal eye discs stained for anti-cleaved *Drosophila* death caspase1 (Dcp-I). (A-D) At the pupal stage, control flies showed basal levels of Dcp-1 positive cells while *Htt93Q* flies have a significantly enhanced level of cell death which is reduced upon downregulation of *Pten*. (E-H) Similarly, *Htt138Q*.*mRFP* flies represent a heightened level of cell death which is significantly reduced by knockdown of *Pten*. (I) Graph showing the average number of Dcp-1 puncta per 100 µm^2^ of the pupal eye disc. (J) The graph represents the average Dcp-1 positive cells in the larval eye disc. Student’s t-test, error bars represent mean± SE (ns= non-significant, ***P<0.001, ****P<0.0001). Red-(A-D) Dcp-1; Blue: DAPI; Green: Dcp-1 (E-H). Scale bar: (A-D) 10µm, (E-H) 50µm.

### Downregulation of Pten restrict cell death in HD disease model

We next investigated whether reduced levels of Pten level alleviate Htt-mediated apoptosis corresponding with the reduction in aggregates. The level of activated death caspase was examined using cleaved Drosophila death caspase 1 (Dcp-1) antibody in the larval and pupal eye disc. The basal level of Dcp-1 staining was observed in the pupal eye disc of control flies (Fig. 5A-B). Increased apoptosis was observed in the pupal eye disc of *Htt93Q* flies as shown by increased Dcp-1 staining (Fig. 5C). Reduction in Pupal eye cell death was demonstrated by a significant reduction in Dcp-1 positive cells in comparison to diseased tissue (arrow in Fig. 5A-D; also see N). The above results were substantiated further in other diseased lines with *Htt138Q* repeats. In comparison to *GMR-Gal4, Htt138Q* expressing eye disc tissue at the larval stage also displayed an enhanced level of Dcp-1 positive puncta (compare Fig. 5E with G; also see J). Notably, the downregulation of *Pten* (compare Fig. 5G-H) caused a significant reduction in the abundance of Dcp-1 positive signal. The above observations demonstrate that the downregulation of *Pten* provides neuroprotection by restricting the Htt-mediated apoptosis in the *Drosophila* model.

### Inhibitor of Ptenimproves the aggregates and behavior

PTEN, as a neuromodulator, can be effectively downregulated using targeted drugs. PTEN inhibitors including VO-OHpic, bpV (Phen), bpV(pic), and SF160 were identified to be specific and effective (Spinelli et al., 2015). VO-OHpic, a potent PTEN inhibitor, has been shown to effectively inhibit Pten in *Drosophila* [37, 43] We sought to identify the optimum concentration of the inhibitor that will inhibit PTEN and reduce the poly(Q) aggregates.

In our dosage screening with *HttQ138*.*mRFP*, 30μM and 50μM inhibitor concentrations were identified to be effective in reducing *HttQ138*.*mRFP* induced aggregates level (Fig. S5A-C). Independent sets of *HttQ138*.*mRFP-expressing* flies at the embryonic stage were allowed to hatch and develop on 30 and 50μM VO-OHic supplemented food. RFP intensity of the 3rd instar larval eye tissues revealed a remarkable reduction in the abundance of poly(Q) aggregates in the groups fed on both 30μM (Fig. S5B) and 50μM PTENi (Fig. S5C, 6B), compared to the DMSO control group (Fig. 6B). These results demonstrated that the identified PTENi administration could efficiently reduce the abundance of poly(Q) aggregates in HD models of *Drosophila*. Moreover, the flies have reduced cell death in the eye disc (Fig. 6B-C, n=10). Furthermore, the climbing ability of flies fed with PTEN inhibitor showed a significant improvement (Fig. 6D, S6. Video).

**Fig. 6.**
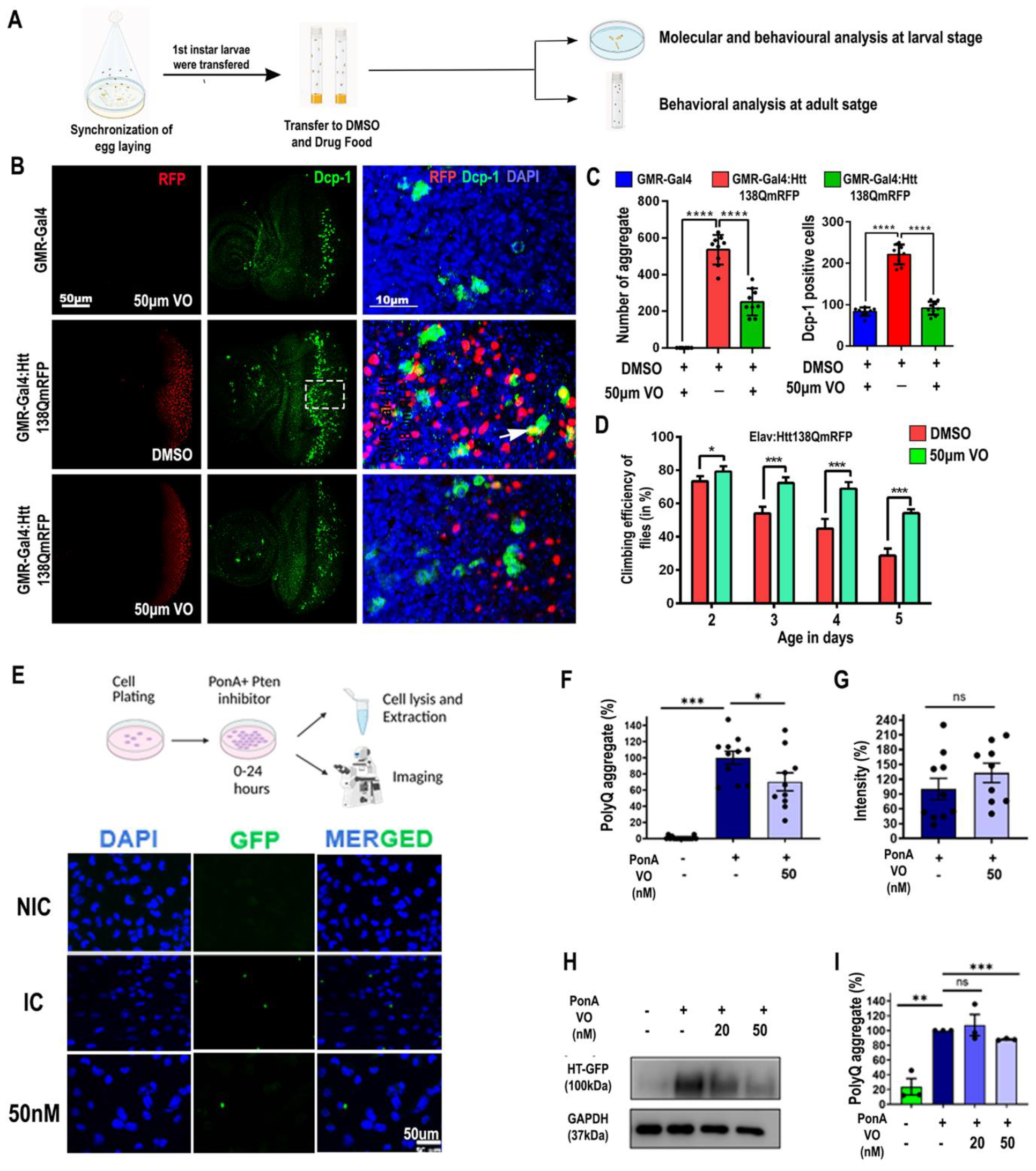
PTEN inhibitor, VO-OHpic (referred to as PTENi), reduces mutant Htt aggregation and cell death. (A) Schematic representation of drug feeding method to Htt expressing flies. (B) 50µM, VO-OHpic treatment reduces Htt aggregation and cell death in compared to DMSO-fed flies. (C) Bar Graph representing the reduction of Htt aggregates and Dcp-I positive cells *in Htt138Q*.*mRFP* files. Student’s t-test, error bars represent mean± SE (ns= non-significant, ****P<0.0001). (D)VO-OHpic flies show improvement in their climbing efficiency compared to DMSO-treated *Elav-Gal4/UAS-Htt138Q*.*mRFP* flies at 2-, 3-, 4-and 5-day aging. One-way ANOVA, error bars represent mean± SE (*P<0.05, ***P<0.001). Green: Dcp-I, Red: *Htt138Q*.*mRFP* and Blue: DAPI. (E) 24-hour Treatment protocol for experiments and at the end of the treatment, cells were scraped for protein isolation and fixed for imaging and representative fluorescence images indicating reduction of poly(Q) aggregates by PTEN inhibitor. (F) Quantification of percentage in the change in Poly(Q) aggregate. (G) Quantification of percentage in the change in intensity. (H) Representatives immunoblot of PTEN inhibitor (w.r.t. IC) and (I) Quantification of the immunoblot showing a reduction in band intensity across the treatment (n=3). (All graphs represented in Mean ± SEM, p values <0.05 considered as statistically significant). Green: Poly(Q) aggregate, Blue: DAPI.

Furthermore, to fortify our findings and to investigate the possible effect of the PTEN inhibitor on lowering the poly(Q) aggregates; we followed the paradigm of co-treating the Mouse neuroblastoma N2a cells with PonA (induction) and VO-OHpic (PTENi) as described in the schematic figure (Fig. 6E, n=3 replicates). We checked for the EGFP puncta per cell as an indirect way of measuring the mHtt aggregates. Similarly, we observed a significant reduction of the puncta per cell in the PTENi-treated conditions, as compared to the inducible control (IC) (only PonA) (Fig. 6F). We also looked for the fluorescence intensity of the EGFP aggregates and the area of the EGFP puncta to understand whether by inhibiting PTEN, there were any significant changes in the intensity of the poly(Q) aggregates (Fig. 6G). We discovered that the VO-OHic-treated cells showed no changes in the intensity and the size of the puncta. To confirm the observation from the fluorescence imaging, we looked for the GFP protein aggregates level to again surrogate measure the protein content of mHtt protein aggregates present in the cell. We found a reduction in the protein aggregates in 50nM of VO-OHic. (Fig.6 H, I). These results suggest that even in cellular conditions the PTEN inhibition may somehow play an active role in altering the mutant aggregate protein level by an unknown mechanism.

## Discussion

Huntington’s disease is a neurodegenerative condition triggered by the abnormal expansion of polyglutamine poly(Q) repeats of the Huntingtin protein. These poly(Q) aggregates instigate widespread cell death within certain brain regions [44]. In animal models, overexpression of mutant poly(Q) proteins causes apoptosis in the eye and brain region [44-46]. Several growth hormone signaling defects have been implicated in Huntington’s disease. For example, mutant Htt protein alters the distribution of EGFR and inhibits ERK signaling [47]. Similarly, in *Drosophila*, expanded poly(Q) repeat prevented EGFR signaling and prevented phosphorylation of ERK [48]. Furthermore, Insulin-like growth factor peptide 1 (IGF1), which activates the insulin signaling pathway by binding to the insulin receptor (a receptor tyrosine kinase), is consistently diminished in animal models and HD patients [49]. Interestingly, in mouse models, intranasal administration of IGF1 has demonstrated the potential to enhance motor function [50]. Furthermore, the administration of Insulin-like growth factor 2 (IGF2) has been shown to reduce poly(Q) aggregates in a mouse model of Huntington’s disease, offering a potential therapeutic avenue [51]. Moreover, administration of FGF has been shown to improve cell survival in Huntington’s disease model [52]. Interestingly, the molecular signaling through EGFR, FGFR, and IGF are regulated by AKT and PTEN is pivotal in the regulation of downstream PI3K/AKT, and loss of its activity causes activation of AKT signaling[53-55]. Since PTEN is a common signaling molecule in the various growth signaling pathways, we hypothesize that PTEN could be a potential avenue for Huntington’s disease therapeutics. In this study, we investigated whether modulating the expression of Pten could ameliorate Huntington’s disease phenotype in the *Drosophila* model system and Huntington’s disease-induced 150Q repeats N2a cell line. Our study demonstrated that the knockdown of Pten improved the rough eye phenotype and loss of photoreceptor neurons caused by overexpression of *Htt93Q* and *Htt138Q* in *Drosophila* models. In addition, knockdown of Pten improved the morphology of the mushroom body which was truncated by mutant Htt proteins. Notably, the morphological improvement was translated into functional improvement as the flies expressing Pten.RNAi with mutant Htt have significantly improved visibility, motor ability, and longevity.

Moreover, pathogenic markers of HD such as accumulation of poly(Q) aggregates were significantly reduced indicating the involvement of the proteasomal or autophagic clearance pathways via Pten downregulation. Hence, these pathways deserve further investigation in subsequent studies. Since Pten is a tumor suppressor, a negative regulator of Pi3K/Akt signaling involved in tissue growth [56], we hypothesize that the knockdown of Pten possibly provided neuroprotection in HD models by reducing the mutant Htt-induced cell death. As expected, the downregulation of Pten reduced the cell death induced by mutant Htt. Our finding from the *Drosophila* model suggests that PTEN downregulation has potential therapeutic application for HD.

Interestingly, studies have demonstrated that the administration of PTEN inhibitors improved neurodegenerative disease models. In the Alzheimer’s disease (AD) mouse model, the administration of PTEN inhibitor, VO-OHpic, improved cognitive deficits, and Long-term potentiation (LTP) in hippocampal neurons [57]. Similarly, inhibition of PTEN provided neuroprotection against axon injury and improved regeneration of axons [58]. Even in the Amyotrophic Lateral Sclerosis disease (ALS) mouse model, inhibition of PTEN improved the survival of motor neurons by improving the neuromuscular junction [59]. With the same rationale, we tested the therapeutic applicability of Pten by feeding the HD fly model with a PTEN inhibitor. In our case, we selected VO-OHpic as a potent inhibitor of PTEN in vivo and in vitro [43]. We administered the VO-OHpic (Pten inhibitor) by incorporating the drug into *Drosophila* food media and allowed the fly to develop. Flies with Huntington’s disease, when fed with the drug-incorporated food, exhibited a notable decrease in aggregate and apoptotic marker levels. Similarly, in the neuronal cell model of HD, VO-OHpic administration significantly reduced the aggregate levels. The reduction of HD markers was also translated into functional improvement as PTEN inhibitor-fed flies exhibited improved climbing ability compared to the flies fed with normal media. Mechanistically, inhibition of PTEN could be altering the defects caused by mutant Huntingtin in PI3K/AKT signaling. Given that AKT signaling serves as a metabolic regulator of Insulin signaling and glucose metabolism pathways, further exploration into the metabolic aspects of this signaling holds promise for future investigations. In HD patients the uptake of glucose is reduced in the brain, and it is interesting to note that metabolic and transcriptomic analysis have demonstrated changes in glucose metabolism and altered gene expression of genes involved in glucose transport [60-62]. Overexpression of the Glucose transporter ameliorated the HD phenotype in the *Drosophila* model [63]. Considering the studies mentioned above, it is highly possible that inhibition of PTEN activated AKT signaling which in turn improved the glucose uptake and suppressed the neuronal cell death. To summarize, our study suggests that Pten is neuroprotective against mutant Htt by reducing the aggregate and ameliorating the survival of neuronal cells (Fig. 7). Our findings highlight the remarkable potential of PTEN inhibition as a promising avenue for advancing therapeutics in Huntington’s disease. Thereby offering renewed hope for those affected by this debilitating condition.

**Fig. 7.**
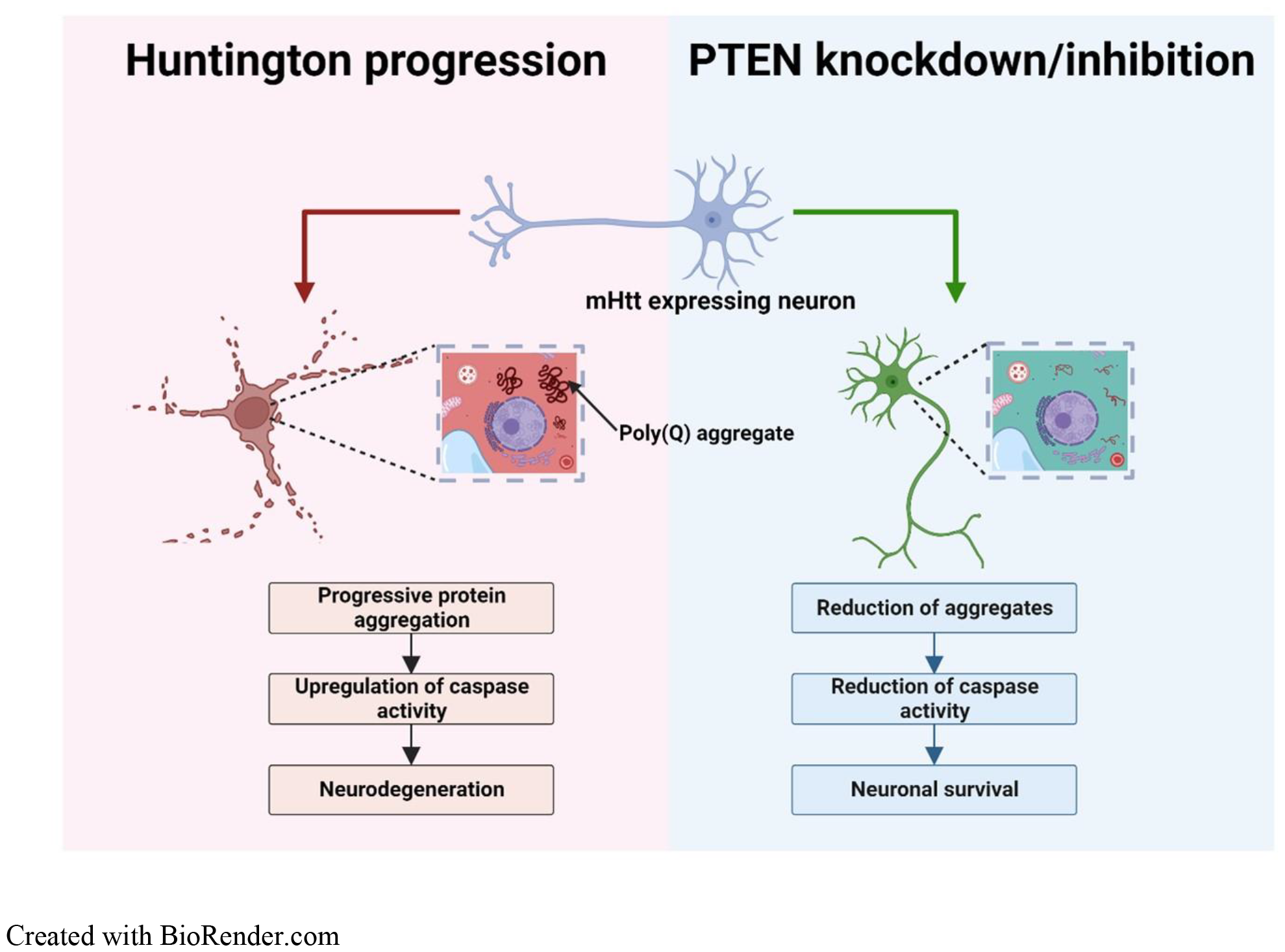
A Schematic model showing how Pten knockdown reduces poly(Q) toxicity. Mutant Huntingtin protein forms poly(Q) aggregates which induces neuronal apoptosis. Reduction of Pten expression restricts aggregate formation, reduces cell death, and promotes the survival of the neuron.

## Acknowledgements

We thank J. Troy Littleton, Massachusetts Institute of Technology, USA for providing the UAS-Htt138Q.mRFP. We thank Bloomington Drosophila Stock Centre (BDSC), USA for the Drosophila stocks. This work was supported by the NBRC Core fund. We thank the NBRC Central Instrument Facility (CIF) for providing the microscopy facility. We thank Mithlesh Kumar Singh and Hari Shankar for their technical help.

## Author contribution

N. and M.D.S. conceptualized and designed the experiments. N., D.T., B.G., A.M., and S.A. carried out the experiments. N., D.T., B.G., A.M., B.S.S. and M.D.S. wrote the manuscript. All the authors discussed the result and approved the final manuscript.

## References

[1] E. M. Gatto, N. G. Rojas, G. Persi, J. L. Etcheverry, M. E. Cesarini, and C. Perandones, ‘Huntington disease: Advances in the understanding of its mechanisms’, Clin Park Relat Disord, vol. 3, p. 100056, 2020, doi: 10.1016/j.prdoa.2020.100056.

[2] S. J. Tabrizi, M. D. Flower, C. A. Ross, and E. J. Wild, ‘Huntington disease: new insights into molecular pathogenesis and therapeutic opportunities’, Nat Rev Neurol, vol. 16, no. 10, pp. 529–546, Oct. 2020, doi: 10.1038/s41582-020-0389-4.

[3] P. McColgan and S. J. Tabrizi, ‘Huntington’s disease: a clinical review’, Eur J Neurol, vol. 25, no. 1, pp. 24–34, Jan. 2018, doi: 10.1111/ene.13413.

[4] A. Palaiogeorgou et al., ‘Recent approaches on Huntington’s disease (Review)’, Biomed Rep, vol. 18, no. 1, p. 5, Nov. 2022, doi: 10.3892/br.2022.1587.

[5] H. S. Bakels, R. A. C. Roos, W. M. C. van Roon-Mom, and S. T. de Bot, ‘Juvenile-Onset Huntington Disease Pathophysiology and Neurodevelopment: A Review’, Movement Disorders, vol. 37, no. 1, pp. 16–24, Jan. 2022, doi: 10.1002/mds.28823.

[6] G. D. Nguyen, S. Gokhan, A. E. Molero, and M. F. Mehler, ‘Selective Roles of Normal and Mutant Huntingtin in Neural Induction and Early Neurogenesis’, PLoS One, vol. 8, no. 5, p. e64368, May 2013, doi: 10.1371/journal.pone.0064368.

[7] M. Jimenez-Sanchez, F. Licitra, B. R. Underwood, and D. C. Rubinsztein, ‘Huntington’s Disease: Mechanisms of Pathogenesis and Therapeutic Strategies’, Cold Spring Harb Perspect Med, vol. 7, no. 7, p. a024240, Jul. 2017, doi: 10.1101/cshperspect.a024240.

[8] C. A. Ross and S. J. Tabrizi, ‘Huntington’s disease: from molecular pathogenesis to clinical treatment’, Lancet Neurol, vol. 10, no. 1, pp. 83–98, Jan. 2011, doi: 10.1016/S1474-4422(10)70245-3.

[9] W. Song, N. Zsindely, A. Faragó, J. L. Marsh, and L. Bodai, ‘Systematic genetic interaction studies identify histone demethylase Utx as potential target for ameliorating Huntington’s disease’, Hum Mol Genet, vol. 27, no. 4, pp. 759–759, Feb. 2018, doi: 10.1093/hmg/ddy020.

[10] L. Pruccoli, C. Breda, G. Teti, M. Falconi, F. Giorgini, and A. Tarozzi, ‘Esculetin Provides Neuroprotection against Mutant Huntingtin-Induced Toxicity in Huntington’s Disease Models’, Pharmaceuticals, vol. 14, no. 10, p. 1044, Oct. 2021, doi: 10.3390/ph14101044.

[11] J. R. Steinert, S. Campesan, P. Richards, C. P. Kyriacou, I. D. Forsythe, and F. Giorgini, ‘Rab11 rescues synaptic dysfunction and behavioural deficits in a Drosophila model of Huntington’s disease’, Hum Mol Genet, vol. 21, no. 13, pp. 2912–2922, Jul. 2012, doi: 10.1093/hmg/dds117.

[12] I. Dragatsis, M. S. Levine, and S. Zeitlin, ‘Inactivation of Hdh in the brain and testis results in progressive neurodegeneration and sterility in mice’, Nat Genet, vol. 26, no. 3, pp. 300–306, Nov. 2000, doi: 10.1038/81593.

[13] C.-Y. Chen, J. Chen, L. He, and B. L. Stiles, ‘PTEN: Tumor Suppressor and Metabolic Regulator’, Front Endocrinol (Lausanne), vol. 9, Jul. 2018, doi: 10.3389/fendo.2018.00338.

[14] S. Dowler et al., ‘Identification of pleckstrin-homology-domain-containing proteins with novel phosphoinositide-binding specificities’, Biochemical Journal, vol. 351, no. 1, p. 19, Oct. 2000, doi: 10.1042/0264-6021:3510019.

[15] Y. Z. Li, A. Di Cristofano, and M. Woo, ‘Metabolic Role of PTEN in Insulin Signaling and Resistance’, Cold Spring Harb Perspect Med, vol. 10, no. 8, p. a036137, Aug. 2020, doi: 10.1101/cshperspect.a036137.

[16] C. Blanco-Aparicio, O. Renner, J. F. M. Leal, and A. Carnero, ‘PTEN, more than the AKT pathway’, Carcinogenesis, vol. 28, no. 7, pp. 1379–1386, Jul. 2007, doi: 10.1093/carcin/bgm052.

[17] D. R. Mattoon, B. Lamothe, I. Lax, and J. Schlessinger, ‘The docking protein Gab1 is the primary mediator of EGF-stimulated activation of the PI-3K/Akt cell survival pathway’, BMC Biol, vol. 2, no. 1, p. 24, 2004, doi: 10.1186/1741-7007-2-24.

[18] Y. He et al., ‘Targeting PI3K/Akt signal transduction for cancer therapy’, Signal Transduct Target Ther, vol. 6, no. 1, p. 425, Dec. 2021, doi: 10.1038/s41392-021-00828-5.

[19] P. Kermer, N. Klöcker, M. Labes, and M. Bähr, ‘Insulin-Like Growth Factor-I Protects Axotomized Rat Retinal Ganglion Cells from Secondary Death via PI3-K-Dependent Akt Phosphorylation and Inhibition of Caspase-3 In Vivo’, The Journal of Neuroscience, vol. 20, no. 2, pp. 722–728, Jan. 2000, doi: 10.1523/JNEUROSCI.20-02-00722.2000.

[20] R. Zoncu, A. Efeyan, and D. M. Sabatini, ‘mTOR: from growth signal integration to cancer, diabetes and ageing’, Nat Rev Mol Cell Biol, vol. 12, no. 1, pp. 21–35, Jan. 2011, doi: 10.1038/nrm3025.

[21] D. C. I. Goberdhan, N. Paricio, E. C. Goodman, M. Mlodzik, and C. Wilson, ‘Drosophila tumor suppressor PTEN controls cell size and number by antagonizing the Chico/PI3-kinase signaling pathway’, Genes Dev, vol. 13, no. 24, pp. 3244–3258, Dec. 1999, doi: 10.1101/gad.13.24.3244.

[22] H. Huang et al., ‘PTEN affects cell size, cell proliferation and apoptosis during Drosophila eye development’, Development, vol. 126, no. 23, pp. 5365–5372, Dec. 1999, doi: 10.1242/dev.126.23.5365.

[23] K. Nowak, G. Seisenbacher, E. Hafen, and H. Stocker, ‘Nutrient restriction enhances the proliferative potential of cells lacking the tumor suppressor PTEN in mitotic tissues’, Elife, vol. 2, Jul. 2013, doi: 10.7554/eLife.00380.

[24] X. Gao, T. P. Neufeld, and D. Pan, ‘Drosophila PTEN Regulates Cell Growth and Proliferation through PI3K-Dependent and -Independent Pathways’, Dev Biol, vol. 221, no. 2, pp. 404–418, May 2000, doi: 10.1006/dbio.2000.9680.

[25] V. K. Godena and K. Ning, ‘Phosphatase and tensin homologue: a therapeutic target for SMA’, Signal Transduct Target Ther, vol. 2, no. 1, p. 17038, Sep. 2017, doi: 10.1038/sigtrans.2017.38.

[26] D. Zhao et al., ‘PTEN Inhibition Protects Against Experimental Intracerebral Hemorrhage-Induced Brain Injury Through PTEN/E2F1/β-Catenin Pathway’, Front Mol Neurosci, vol. 12, Dec. 2019, doi: 10.3389/fnmol.2019.00281.

[27] K. R. Weiss, Y. Kimura, W.-C. M. Lee, and J. T. Littleton, ‘Huntingtin Aggregation Kinetics and Their Pathological Role in a Drosophila Huntington’s Disease Model’, Genetics, vol. 190, no. 2, pp. 581–600, Feb. 2012, doi: 10.1534/genetics.111.133710.

[28] J. S. Steffan et al., ‘Histone deacetylase inhibitors arrest polyglutamine-dependent neurodegeneration in Drosophila’, Nature, vol. 413, no. 6857, pp. 739–743, Oct. 2001, doi: 10.1038/35099568.

[29] B. A. Hay, T. Wolff, and G. M. Rubin, ‘Expression of baculovirus P35 prevents cell death in Drosophila’, Development, vol. 120, no. 8, pp. 2121–2129, Aug. 1994, doi: 10.1242/dev.120.8.2121.

[30] J. M. Sidisky and D. T. Babcock, ‘Visualizing Synaptic Degeneration in Adult *Drosophila* in Association with Neurodegeneration’, Journal of Visualized Experiments, no. 159, May 2020, doi: 10.3791/61363.

[31] J. Schindelin et al., ‘Fiji: an open-source platform for biological-image analysis’, Nat Methods, vol. 9, no. 7, pp. 676–682, Jul. 2012, doi: 10.1038/nmeth.2019.

[32] C. Arshadi, U. Günther, M. Eddison, K. I. S. Harrington, and T. A. Ferreira, ‘SNT: a unifying toolbox for quantification of neuronal anatomy’, Nat Methods, vol. 18, no. 4, pp. 374–377, Apr. 2021, doi: 10.1038/s41592-021-01105-7.

[33] M. H. Longair, D. A. Baker, and J. D. Armstrong, ‘Simple Neurite Tracer: open source software for reconstruction, visualization and analysis of neuronal processes’, Bioinformatics, vol. 27, no. 17, pp. 2453–2454, Sep. 2011, doi: 10.1093/bioinformatics/btr390.

[34] J. M. Sidisky, D. Weaver, S. Hussain, M. Okumus, R. Caratenuto, and D. Babcock, ‘Mayday sustains trans-synaptic BMP signaling required for synaptic maintenance with age’, Elife, vol. 10, Mar. 2021, doi: 10.7554/eLife.54932.

[35] Nisha and S. Sarkar, ‘Downregulation of glob1 suppresses pathogenesis of human neuronal tauopathies in Drosophila by regulating tau phosphorylation and ROS generation’, Neurochem Int, vol. 146, p. 105040, Jun. 2021, doi: 10.1016/j.neuint.2021.105040.

[36] M. D. Singh et al., ‘NCBP2 modulates neurodevelopmental defects of the 3q29 deletion in Drosophila and Xenopus laevis models’, PLoS Genet, vol. 16, no. 2, p. e1008590, Feb. 2020, doi: 10.1371/journal.pgen.1008590.

[37] J. Lattner, W. Leng, E. Knust, M. Brankatschk, and D. Flores-Benitez, ‘Crumbs organizes the transport machinery by regulating apical levels of PI(4,5)P2 in Drosophila’, Elife, vol. 8, Nov. 2019, doi: 10.7554/eLife.50900.

[38] S. A. Deshpande et al., ‘Quantifying Drosophila food intake: Comparative analysis of current methodology’, Nat Methods, vol. 11, no. 5, pp. 535–540, 2014, doi: 10.1038/nmeth.2899.

[39] G. Roman and R. L. Davis, ‘Molecular biology and anatomy of Drosophila olfactory associative learning’, BioEssays, vol. 23, no. 7, pp. 571–581, Jul. 2001, doi: 10.1002/bies.1083.

[40] R. L. Davis, ‘Mushroom Bodies, Ca2+ Oscillations, and the Memory Gene amnesiac’, Neuron, vol. 30, no. 3, pp. 653–656, May 2001, doi: 10.1016/S0896-6273(01)00329-4.

[41] F. Saudou and S. Humbert, ‘The Biology of Huntingtin’, Neuron, vol. 89, no. 5, pp. 910–926, Mar. 2016, doi: 10.1016/j.neuron.2016.02.003.

[42] A. Couto, M. Alenius, and B. J. Dickson, ‘Molecular, Anatomical, and Functional Organization of the Drosophila Olfactory System’, Current Biology, vol. 15, no. 17, pp. 1535–1547, Sep. 2005, doi: 10.1016/j.cub.2005.07.034.

[43] L. H. Mak, R. Vilar, and R. Woscholski, ‘Characterisation of the PTEN inhibitor VO-OHpic’, J Chem Biol, vol. 3, no. 4, pp. 157–163, Oct. 2010, doi: 10.1007/s12154-010-0041-7.

[44] C. Portera-Cailliau, J. Hedreen, D. Price, and V. Koliatsos, ‘Evidence for apoptotic cell death in Huntington disease and excitotoxic animal models’, The Journal of Neuroscience, vol. 15, no. 5, pp. 3775–3787, May 1995, doi: 10.1523/JNEUROSCI.15-05-03775.1995.

[45] F. Saudou, S. Finkbeiner, D. Devys, and M. E. Greenberg, ‘Huntingtin Acts in the Nucleus to Induce Apoptosis but Death Does Not Correlate with the Formation of Intranuclear Inclusions’, Cell, vol. 95, no. 1, pp. 55–66, Oct. 1998, doi: 10.1016/S0092-8674(00)81782-1.

[46] P. W. Faber, J. R. Alter, M. E. MacDonald, and A. C. Hart, ‘Polyglutamine-mediated dysfunction and apoptotic death of a Caenorhabditis elegans sensory neuron’, Proceedings of the National Academy of Sciences, vol. 96, no. 1, pp. 179–184, Jan. 1999, doi: 10.1073/pnas.96.1.179.

[47] M. A. B. Melone et al., ‘Mutant huntingtin regulates EGF receptor fate in non-neuronal cells lacking wild-type protein’, Biochimica et Biophysica Acta (BBA) - Molecular Basis of Disease, vol. 1832, no. 1, pp. 105–113, Jan. 2013, doi: 10.1016/j.bbadis.2012.09.001.

[48] J.-C. Liévens, T. Rival, M. Iché, H. Chneiweiss, and S. Birman, ‘Expanded polyglutamine peptides disrupt EGF receptor signaling and glutamate transporter expression in Drosophila’, Hum Mol Genet, vol. 14, no. 5, pp. 713–724, Mar. 2005, doi: 10.1093/hmg/ddi067.

[49] M. A. Pouladi et al., ‘Full-length huntingtin levels modulate body weight by influencing insulin-like growth factor 1 expression’, Hum Mol Genet, vol. 19, no. 8, pp. 1528–1538, Apr. 2010, doi: 10.1093/hmg/ddq026.

[50] C. Lopes et al., ‘IGF-1 Intranasal Administration Rescues Huntington’s Disease Phenotypes in YAC128 Mice’, Mol Neurobiol, vol. 49, no. 3, pp. 1126–1142, Jun. 2014, doi: 10.1007/s12035-013-8585-5.

[51] P. García-Huerta et al., ‘Insulin-like growth factor 2 (IGF2) protects against Huntington’s disease through the extracellular disposal of protein aggregates’, Acta Neuropathol, vol. 140, no. 5, pp. 737–764, Nov. 2020, doi: 10.1007/s00401-020-02183-1.

[52] A. R. La Spada, ‘Huntington’s disease and neurogenesis: FGF-2 to the rescue?’, Proceedings of the National Academy of Sciences, vol. 102, no. 50, pp. 17889–17890, Dec. 2005, doi: 10.1073/pnas.0509222102.

[53] B. R. Chaffee et al., ‘FGFR and PTEN signaling interact during lens development to regulate cell survival’, Dev Biol, vol. 410, no. 2, pp. 150–163, Feb. 2016, doi: 10.1016/j.ydbio.2015.12.027.

[54] S. R. Shinde and S. Maddika, ‘PTEN modulates EGFR late endocytic trafficking and degradation by dephosphorylating Rab7’, Nat Commun, vol. 7, no. 1, p. 10689, Feb. 2016, doi: 10.1038/ncomms10689.

[55] R. Endersby and S. J. Baker, ‘PTEN signaling in brain: neuropathology and tumorigenesis’, Oncogene, vol. 27, no. 41, pp. 5416–5430, Sep. 2008, doi: 10.1038/onc.2008.239.

[56] J. Liu et al., ‘Insulin activates the insulin receptor to downregulate the PTEN tumour suppressor’, Oncogene, vol. 33, no. 29, pp. 3878–3885, Jul. 2014, doi: 10.1038/onc.2013.347.

[57] S. Frere and I. Slutsky, ‘Targeting PTEN interactions for Alzheimer’s disease’, Nat Neurosci, vol. 19, no. 3, pp. 416–418, Mar. 2016, doi: 10.1038/nn.4248.

[58] Y. Ohtake, U. Hayat, and S. Li, ‘PTEN inhibition and axon regeneration and neural repair’, Neural Regen Res, vol. 10, no. 9, p. 1363, 2015, doi: 10.4103/1673-5374.165496.

[59] J. Wang, L. Tierney, R. Mann, T. Lonsway, and C. L. Walker, ‘Bisperoxovanadium promotes motor neuron survival and neuromuscular innervation in amyotrophic lateral sclerosis’, Mol Brain, vol. 14, no. 1, p. 155, Dec. 2021, doi: 10.1186/s13041-021-00867-7.

[60] G. Chaves et al., ‘Metabolic and transcriptomic analysis of Huntington’s disease model reveal changes in intracellular glucose levels and related genes’, Heliyon, vol. 3, no. 8, p. e00381, Aug. 2017, doi: 10.1016/j.heliyon.2017.e00381.

[61] V. Morea et al., ‘Glucose transportation in the brain and its impairment in Huntington disease: one more shade of the energetic metabolism failure?’, Amino Acids, vol. 49, no. 7, pp. 1147–1157, Jul. 2017, doi: 10.1007/s00726-017-2417-2.

[62] A. Ciarmiello, G. Giovacchini, S. Orobello, L. Bruselli, F. Elifani, and F. Squitieri, ‘18F-FDG PET uptake in the pre-Huntington disease caudate affects the time-to-onset independently of CAG expansion size’, Eur J Nucl Med Mol Imaging, vol. 39, no. 6, pp. 1030–1036, Jun. 2012, doi: 10.1007/s00259-012-2114-z.

[63] M. T. Besson, K. Alegría, P. Garrido-Gerter, L. F. Barros, and J.-C. Liévens, ‘Enhanced Neuronal Glucose Transporter Expression Reveals Metabolic Choice in a HD Drosophila Model’, PLoS One, vol. 10, no. 3, p. e0118765, Mar. 2015, doi: 10.1371/journal.pone.0118765.

