## Supplementary for "Protective role of *Pten* downregulation in Huntington’s Disease models"

Supplementary figures

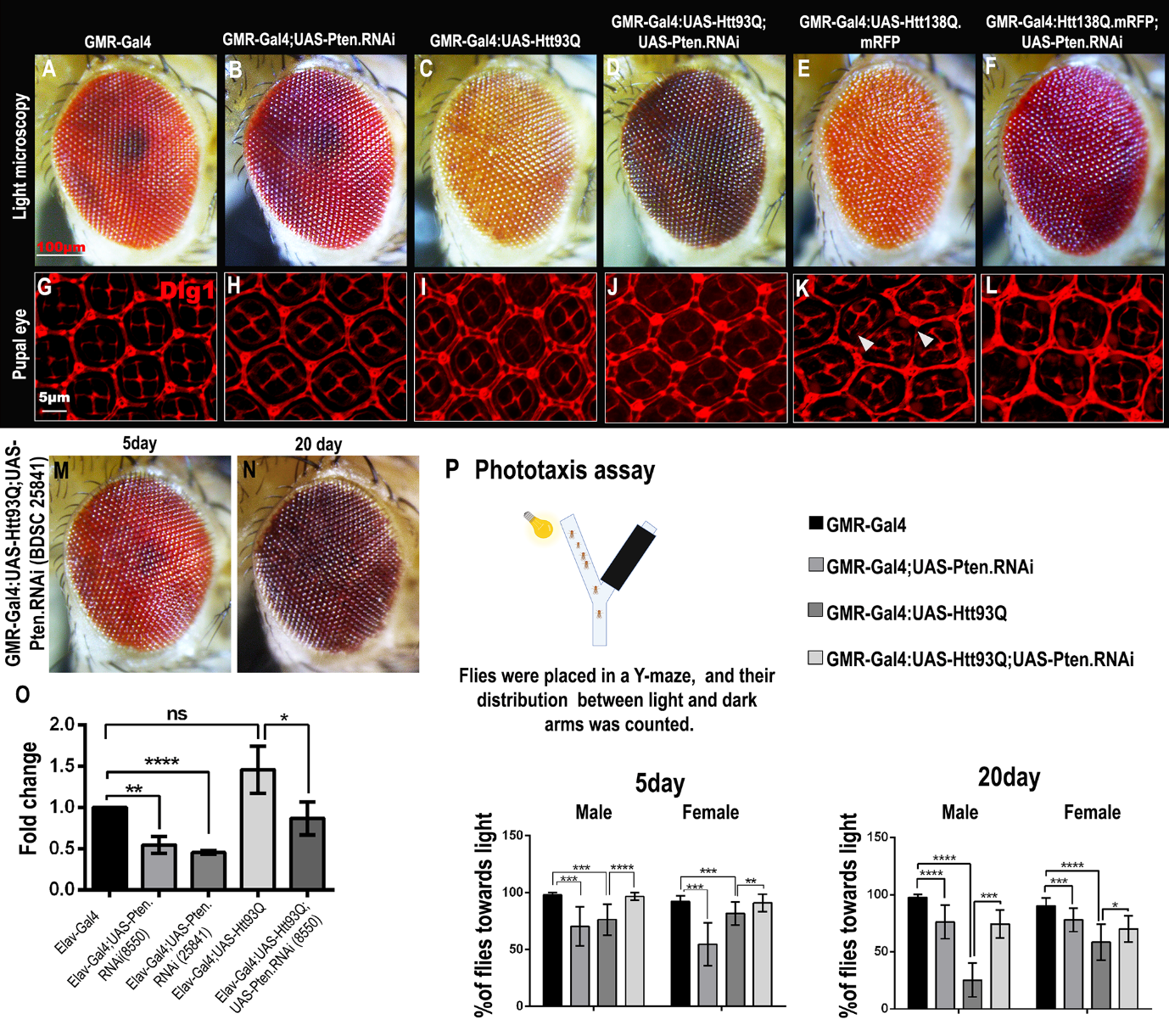

**Figure S1:** **Tissue-specific downregulation of Pten suppresses Htt-induced phenotypes.** (A-F) Bright-field images of 5-day old adult *Drosophila* eyes. (A-B) control flies (C-D) *Htt93Q* flies showed mild depigmentation and reduced eye size which was restored back by downregulation of *Pten*. (E) Similarly, expression of *Htt138Q* exhibited ommatidial fusion and reduced eye size. (F) *Pten.RNAi* improved eye phenotypes. (G-L) 40 hours-old Pupal eye stained with Dlg-1 antibody. (G-H) Control flies. (I) Expression of *Htt93Q* did not show defects in pupal eye. (K-L) *Htt138Q* expression caused structural defects (arrow), which was restored partially by *Pten.RNAi*. (M-N) Second *Pten.RNAi (BDSC#25841)* also suppressed the *Htt93Q* eye phenotype. (O) Real-time PCR data showing downregulation of *Pten* level in control and diseased flies. (P) Schematic representation of phototaxis assay at adult flies by using Y-maze. The bar graph shows the percentage of flies in illuminated arm against the dark arm at 5 day and 20 days old flies. (Student’s t-test ns=not significant, *P<0.05, **P<0.01, ***P<0.001, ****P<0.0001). Scale bars: (A-F) 100μm, (G-L) 5μm.

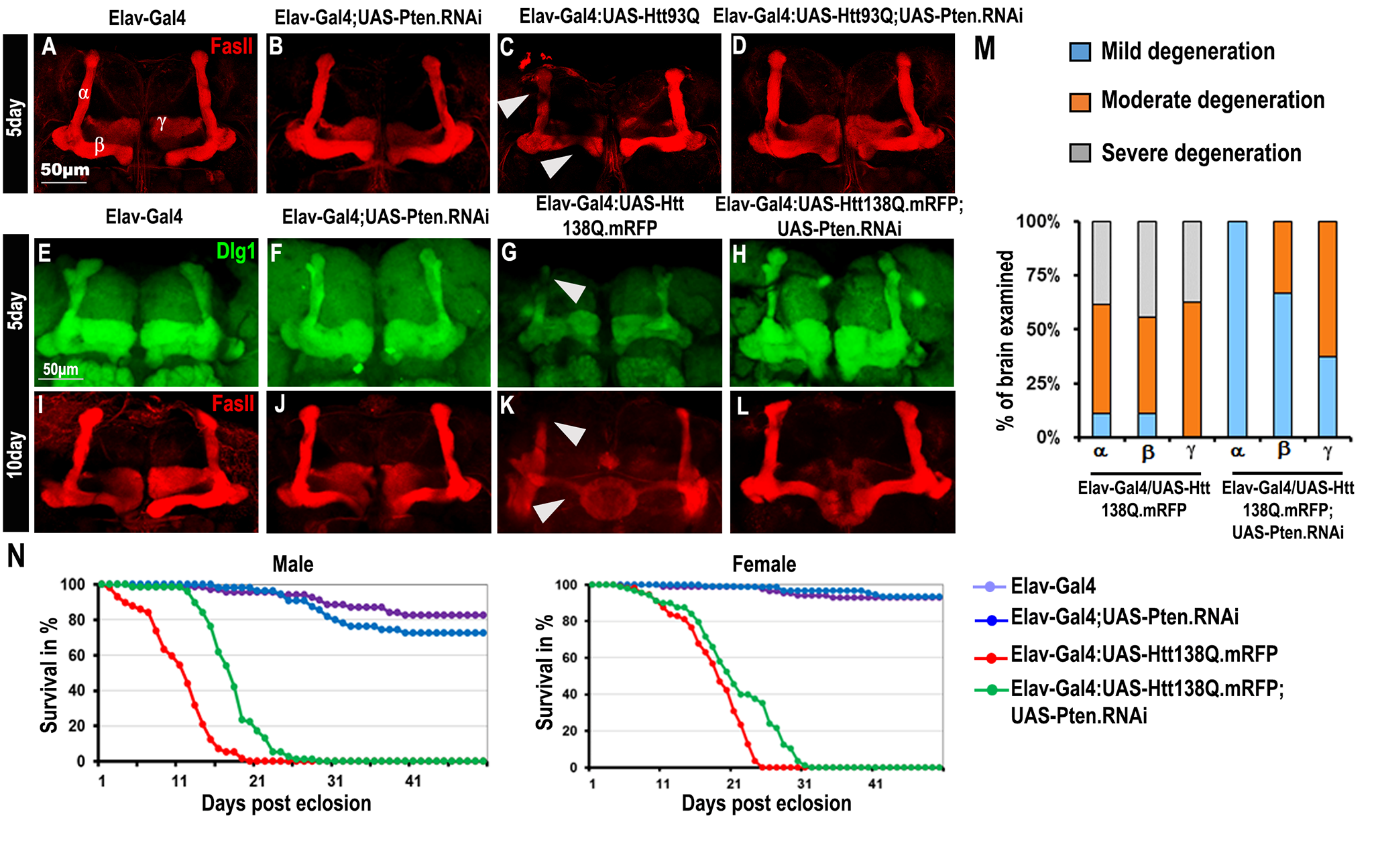

**Figure S2:** **Structural and behavioral improvement by Pten knockdown in the Brain.** (A-D) 5 days-old *Drosophila* adult brain stained with Fas-II antibody. (A, B) Control flies showing α, β, and γ lobes of the mushroom body. (C) Degeneration of α, and β lobe in *Htt93Q* expressing flies. (D) Downregulation of *Pten* restored all three α, β, and γ lobes. (E-F) Control flies. (G) Expression of *Htt138Q.mRFP* caused degeneration of mushroom body structures at 5 day. (H) Pten knockdown in disease background improved the mushroom body structure. (I- J) Control flies. (K-L) *Htt138Q*-induced degeneration of mushroom body was improved by knockdown of Pten in 10-day-old flies. (M) Graphical representation for the percentage of degeneration of α, β, and γ lobes. (N) Survival curve indicating increased longevity when *Pten* was downregulated in *Htt138Q* background. Survival curves were generated by the Kaplan-Meier method and statistical significance was determined by the log-rank test (P<0.05).

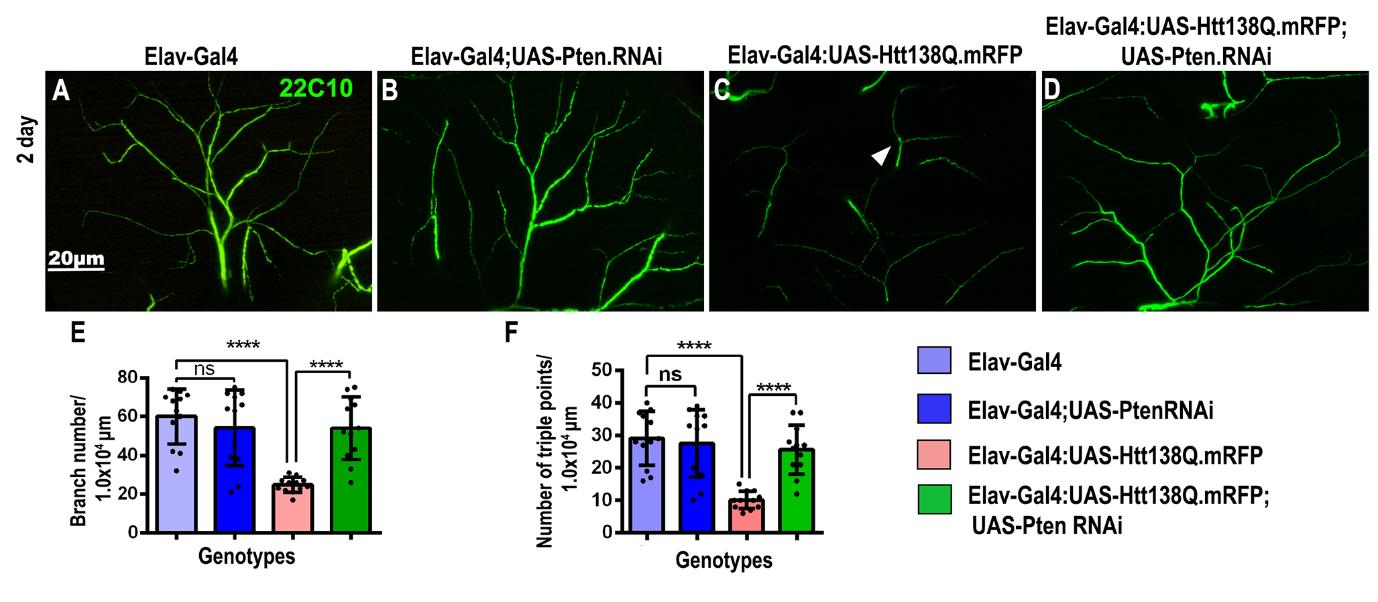

**Figure S3: Pten Knockdown improves the NMJ structure of *Htt138Q* flies.** (A-B**)** Control flies showing well-arranged branching of motor neurons in the Dorso-longitudinal muscles (DLMs). (C) Expression of *Htt138Q* resulted in degeneration of the branching pattern of motor neurons. (D) *Pten* knockdown improved the branching of NMJ in *Htt138Q* flies (arrows represent synaptic degeneration). (E-F) Bar graph showing the quantification of branching number as well as triple points in fixed ROI area in each sample. Student’s t-test, error bars represent mean± SE (, ns=not significant, **P<0.01, ***P<0.001, ****P<0.0001). Green: anti-Futsch (22C10); Scale bar: 20µm.

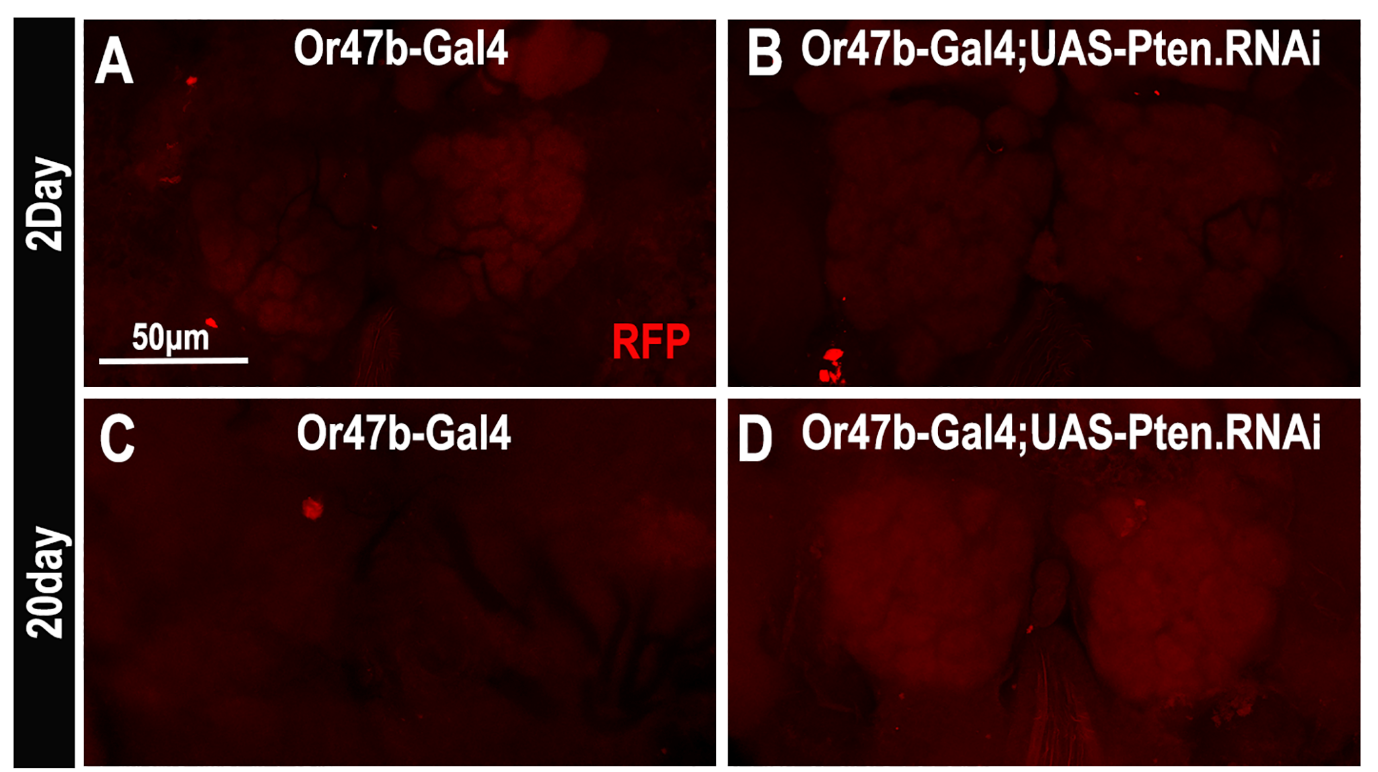

F**igure S4. Or47b-Gal4 expression shows no aggregate in control flies.** (A) 2-day old *Or47b-Gal4* and (B) *Or47b-Gal4/UAS-Pten.RNAi* control flies showed no aggregates. (C-D) A similar observation has been noted in 20-day-old flies also in both *Or47b-Gal4* and *Or47b-Gal4/UAS-Pten.RNAi*. Scale bar: 50µm.

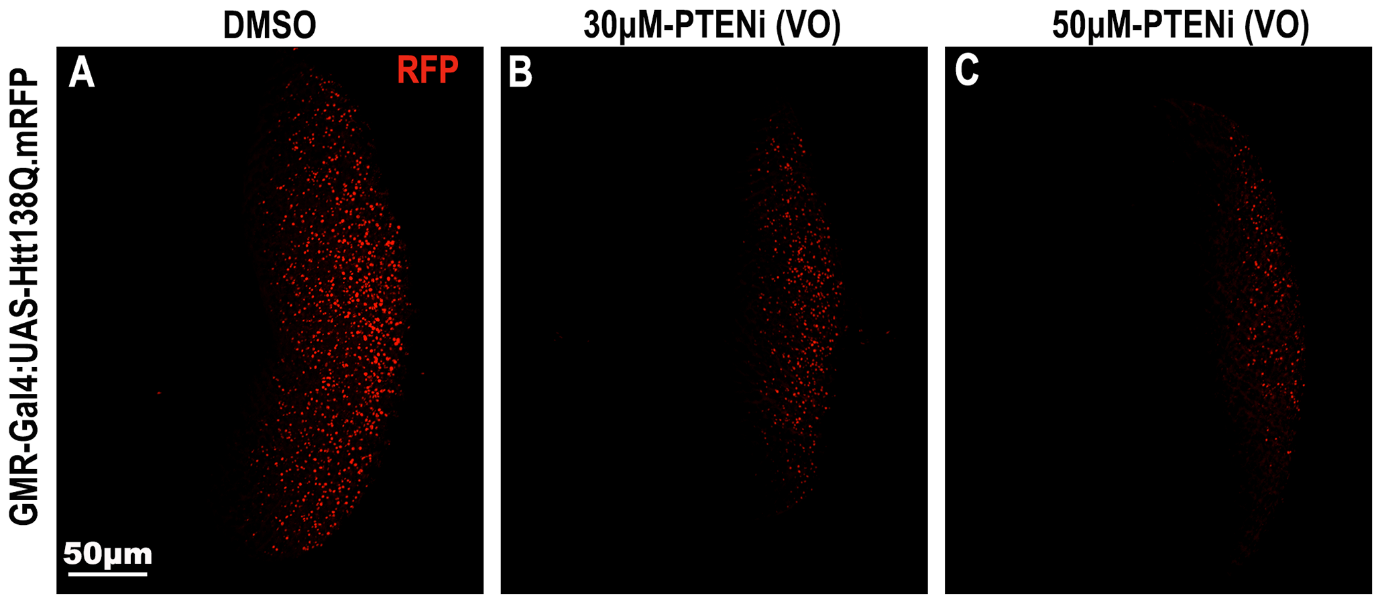

**Figure S5. VO-OHpic (PTENi) reduces poly(Q) aggregate in a dose-dependent manner.** Compared to DMSO-fed flies, Pten inhibitor (VO-OHpic) fed larval eye disc of *GMR-Gal4;UAS-Htt138Q.mRFP* exhibited a significant reduction in the amount of poly(Q) aggregates (Red) with 30m and 50m concentration respectively.

**S6:** Video of climbing efficiency of PTEN inhibitor fed flies at 5-Day.

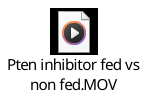
